# Dissecting the Cellular Landscape and Transcriptome Network in Viral Myocarditis by Single-Cell RNA Sequencing

**DOI:** 10.1101/2021.05.16.444367

**Authors:** Ninaad Lasrado, Nicholas Borcherding, Rajkumar Arumugam, Timothy K. Starr, Jay Reddy

## Abstract

Myocarditis induced with Coxsackievirus B3 (CVB3) is commonly employed to study viral pathogenesis in mice. Although infectious virus is cleared after the acute phase, affected animals chronically develop the features of dilated cardiomyopathy, which may involve the mediation of immune and non-immune cells. To dissect this complexity, we performed single-cell RNA sequencing on heart cells obtained from healthy and myocarditic mice, leading us to note that myocarditic mice had significantly higher proportions of myeloid cells, CD4 and CD8 T cells, and fibroblasts, whereas NK cells, ILCs and B cells were low. While the transcriptome profiles of myeloid cells revealed detection of monocytes and macrophages of M2 phenotype with pathways important in immune metabolism and inflammation, T cells consisted of Th17 cells, CTLs, and Treg cells with transcriptome signatures critical for cytotoxic functions. Although fibroblasts detected in myocarditic mice were phenotypically heterogeneous, their transcriptomes played roles in fibrosis and regulation of inflammation and immune responses. Additionally, analysis of intercellular communication networks revealed unique interactions and signaling pathways in the cardiac cellulome, whereas myeloid cells and T cells in myocarditic mice revealed uniquely upregulated transcription factors modulating cardiac remodeling functions. Taken together, our data suggest that M2 cells, T cells, and fibroblasts may cooperatively or independently participate in the pathogenesis of viral myocarditis.

## Introduction

Myocarditis is a significant clinical entity in young infants and adolescents ^1, 2^. While the disease is spontaneously resolved in most affected individuals, ∼20% of those affected develop chronic myocarditis that can lead to dilated cardiomyopathy (DCM)^3^. More recently, the term myocarditis has been designated as inflammatory cardiomyopathy to describe the occurrence of myocarditis in association with cardiac dysfunction ^2^. Furthermore, it is not uncommon to detect low grade inflammation in the hearts of healthy individuals, as has been suggested by a study involving accidental deaths in which heart infiltrates were detected in ∼1 to 9% of autopsies ^4^. Consistent with this finding, the presence of heart infiltrates in the sudden deaths of young athletes have raised a question as to the underlying mechanisms ^5^. Therapeutically, due to a lack of effective treatment options, ∼50% of DCM patients undergo heart transplantation, and children with acute myocarditis only have a ∼60% likelihood of transplantation-free survival ^6^. This is complicated by the finding that myocarditis can result from multiple triggers, whose disease-inducing abilities are complex in nature.

Viruses are the major causative agents of myocarditis^2, 7, 8^. Since it is difficult to study the pathogenic mechanisms of viral myocarditis in humans, animal models are commonly employed. However, infection models for all viruses are not available or feasible for routine experimentation – except for enteroviruses that include B group Coxsackieviruses. Thus, Coxsackievirus B3 (CVB3)-induced myocarditis is commonly employed to investigate the pathogenic mechanisms of viral myocarditis.

Of various rodent species, mice are highly susceptible to CVB3 infection^9^, and their MHC haplotypes influence the disease outcome. While H-2 (IA^b^)-bearing C57Bl/6 mice develop acute infection and are resistant to the development of chronic myocarditis, A/J (IA^k^) and Balb/c (IA^d^) mice develop chronic disease, making them suitable to study the pathogenic mechanisms that may involve both viral and host factors ^7, 9, 10^. We routinely use A/J mice that are highly susceptible to a human isolate of the Nancy strain of CVB3 ^9, 11^. Upon infection, animals develop myocarditis in two phases in continuum: acute or viral phase lasting approximately 14 to 18 days, followed by a chronic or non-viral phase in which cardiac dysfunctional features are manifested ^7^. The chronic nature of CVB3 is particularly relevant to humans because signatures of enteroviruses have been identified in DCM patients, as indicated by the detection of virus-reactive antibodies and the viral genomic material ^2, 8, 12^. These findings raise the question whether residual virus, if any, or reactivation of viral nucleic acid, if at all possible, may potentially contribute to DCM pathogenesis. But conclusive evidence is lacking to support these notions. Another possibility is autoimmunity, as autoantibodies have been detected in both DCM patients and CVB3 infection models in mice ^2, 8, 12–14^. In our studies, by creating MHC class II dextramers for various cardiac antigens, we have demonstrated that CVB3 infection leads to the generation of pathogenic autoreactive T cells with multiple antigen-specificities localized in both lymphoid and non-lymphoid organs with the potential for them to be recirculated back into the heart under inflammatory conditions ^11, 15^.

Nonetheless, if autoimmunity is a key underlying mechanism for DCM pathogenesis, affected patients should be responsive to immune therapies, but mixed successes have been achieved in clinical trials ^3^. Furthermore, the heart is not an immunologically privileged organ, and immune cells have free access to the heart muscle. From the standpoint of immune defense mechanisms, arrival of immune cells to damaged cardiac tissue is expected to be beneficial to the host, but their detrimental effects cannot be discounted if their functions are dysregulated. Additionally, numerous cardiac resident cells ‒ specifically, cardiomyocytes, fibroblasts, and smooth muscle cells ‒ and tissue-resident immune cells, such as macrophages and dendritic cells (DCs), may be severely affected by inflammatory responses, leading to alterations in cardiac functions. Collectively, many cell types may potentially participate in the cardiac dysfunction that culminates in cardiac remodeling events. To dissect this complexity, we used single-cell-RNA sequencing (scRNAseq) to define the cardiac cellulome and its transcriptome profiles during the post-infectious phase of myocarditis in A/J mice. The data revealed detection of mainly myeloid cells, T cells and fibroblasts in the heart infiltrates from myocarditic mice, which may have a role in the development of chronic myocarditis and DCM.

## Results

Using the mouse model of viral myocarditis induced with CVB3, we analyzed cellular infiltrates in hearts to identify novel genes, transcription factors (TFs), and signaling pathways that contribute to disease progression in the viral pathogenesis.

### Myeloid cells, T cells and fibroblasts are the major enriched cell types in hearts of myocarditic mice

To elucidate cellular compositions and diversity in their transcriptome profiles, we performed scRNAseq using heart cells from both mice infected with CVB3 and healthy mice (**Fig 1A**). Single cell suspensions obtained from myocarditic and healthy mice were stained with annexin-V and propidium iodide (PI) for sorting viable cells (annexin-V^-^PI^-^) by flow cytometry (**Fig 1A and Fig S1**). After confirming the viability (100%), 16,000 cells each from the healthy and myocarditic groups were loaded into the 10x genomics chromium 3’expression system, and their libraries were sequenced for downstream analysis (**Fig 1A**).

**Fig 1:**
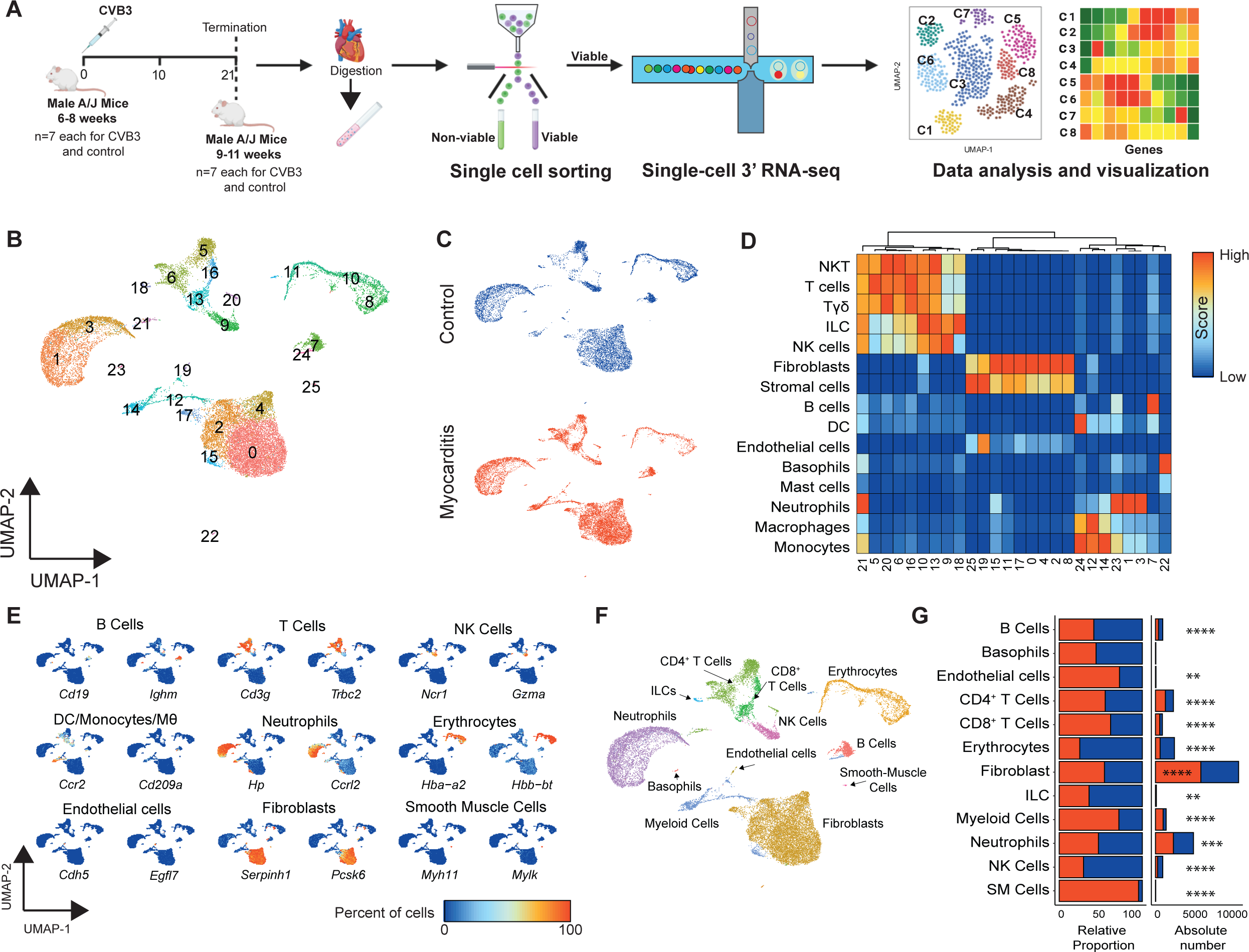
Phenotypic characterization of heart infiltrates in CVB3-infected mice. A. Schematic representation of the experimental approach. Hearts harvested from A/J mice infected with or without CVB3 on day 21 post-infection were enzymatically digested to obtain single-cell suspensions. After sorting the viable cells by flow cytometry as described in Methods, single-cell libraries were prepared and sequenced. Raw data were subjected for downstream analysis to characterize cellular distributions. B. Mapping of cell clusters. Uniform Manifold Approximation and Projection (UMAP) visualization of cells from healthy (9,734 cells) and myocarditic mice (13,251 cells) using Seurat identified 26 different clusters after unsupervised clustering. C. Distribution of cells between treatment groups. UMAP projections showing relative distribution of cell clusters in healthy and myocarditic mice. D. Prediction of cell types. Cell types were predicted using singleR R package ^92^; normalized correlation values for predicted immune cell phenotypes are shown. Cluster of columns based on Ward.D2 distance between normalized correlation values across all pure immune cell populations in the Immgen database ^17^. E. Identification of cell types using select lineage markers. Using the canonical markers, major cell types in both healthy and myocarditic mice were assigned. F. Annotating cell types. Twelve major cell types were identified and annotated based on the expression pattern of canonical cell markers. G. Relative distribution of cell types. Relative distribution of cell types scaled by total number of cells per condition is shown. Red indicates myocarditis, and blue indicates control. Significance best on two-proportion Z-tests with p-values correct for multiple comparisons using the Benjamini-Hochberg method; * < 0.05, ** < 0.01, *** < 0.001, and **** < 0.0001.

A combined total of 22,985 cells from healthy (n=9,734) and myocarditic mice (n=13,251) were analyzed using the Seurat R package ^16^, and an unbiased clustering yielded 26 cell clusters (**Fig 1B**) along a single uniform manifold approximation and projection (UMAP). In general, cells from both control and myocarditic hearts were present in most clusters (**Fig 1C**). We next annotated each cluster by using two approaches: 1) correlation of mouse gene signatures in the Immune Genome Project (ImmGen) database ^17^, and 2) expression of canonical markers, with average gene expression in the clusters. We used the SingleR R package ^18^ to assign cell types to each cluster based on correlations between gene expression in the cluster and gene expression in purified mouse cell populations in the ImmGen database. **Fig 1D** shows the predicted proportions of cells identified as natural killer (NK)T cells, T cells, Tγδ cells, innate lymphoid cells (ILCs), NK cells, fibroblasts, stromal cells, neutrophils, macrophages, and monocytes that were present in multiple clusters, whereas B cells, DCs, endothelial cells (ECs), and basophils were restricted to one cluster each. Next, using canonical markers for indicated cell types (**Table S1**), as shown with two examples for each cell type (**Fig 1E**), we noted that T cells, fibroblasts, neutrophils, and erythroid cells were present at relatively higher proportions than B cells, NK cells, myeloid cells, ECs, and smooth muscle cells (SMCs). Based on the above two approaches, cluster annotations were then made (**Fig 1F**). By comparing the relative proportions of each cell type in both groups, we found that T cells (CD4 and CD8), fibroblasts, and myeloid cells were significantly enriched in myocarditis, whereas neutrophils, NK cells, ILCs, and B cells were reduced in myocarditic mice as compared to healthy mice (**Fig 1G, Table S2**). Although ECs and smooth muscle cells (SMCs) were also elevated in myocarditic mice (**Fig 1G**), their absolute number was relatively low (**Table S2**). Nonetheless, detection in myocarditic mice of the predominant populations of cells described above raised questions as to their significance in CVB3 pathogenesis.

### Transcriptome analysis of myeloid cells led to the detection of predominantly monocytes and macrophages with pathways involved in immune metabolism and inflammation

To understand the contribution of myeloid cells in post-infectious myocarditis, we re-clustered the myeloid cells separately. As shown in **Fig 1B**, cells of the myeloid lineage were scattered in five clusters ‒ 12, 14, 15, 19, and 25. Further unbiased subclustering led us to identify six distinct subpopulations (**Fig 2A**), with cellular proportions varying between healthy and myocarditic mice (**Fig 2B, left panel**). Cells in clusters 0 and 1 were significantly elevated, followed by clusters 4 and 2 in myocarditic mice (**Fig 2B, right panel**). By using canonical myeloid cell markers, cytokines, chemokines, and other molecules as shown (**Fig 2C, Fig S2A**), we identified monocytes, macrophages, cDC1, moDC, and a discrete subset of cells that expressed predominantly mitochondrial genes (Mt-high) (**Fig 2D, left panel**). Although DCs contained two subpopulations, no differential gene expression (DGE) was noted in their transcriptomes. Furthermore, we used Slingshot ^19^ to infer the cell lineage and pseudo-time trajectory, which indicated branching of monocytes into other cell types (**Fig 2D, left panel**). The top immune-related genes that were used to construct this model are shown (**Fig 2D, right panel**), with *Msrb1,* an anti-inflammatory selenoprotein ^20^, making the largest contribution to this developmental trajectory. Evaluation of cell cycle phases revealed monocytes from healthy mice to be in the G2M and/or G1 growth phases, whereas monocytes, macrophages, and moDCs from myocarditic mice were in the S and G1 phases, suggesting that the growth stages of the latter cells might represent their activation status (**Fig 2E**).

**Fig 2:**
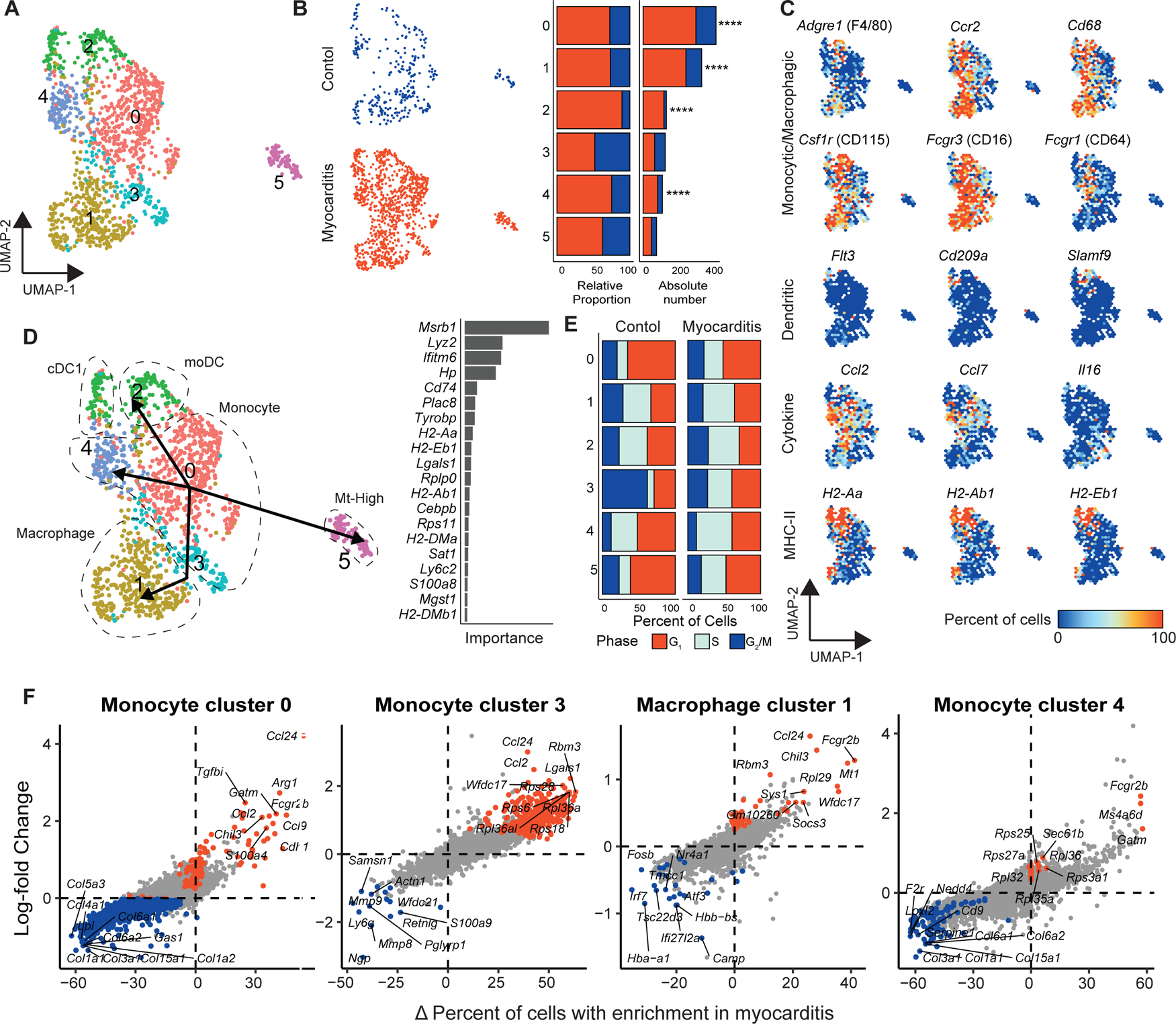
Distribution and characterization of myeloid cells in heart infiltrates. UMAP of myeloid cells identified in heart infiltrates (**A**) and their distributions are shown in healthy controls and myocarditic mice (**B**); relative proportions of cells are indicated by cluster in the bar plot, with red indicating myocarditis, and blue indicating control. By using select markers for various cell types (**C**), individual subsets were then identified (**D**); the cell trajectory is shown using the slingshot method with Cluster 0 as the origin. While the ranked bar graph indicates the top immune-related genes across conditions, the bar graph represents cell cycle phases by cluster between healthy and myocarditic mice (**E**). Percentage difference (Δ Percent of Cells) and log-fold change based on the Wilcoxon rank sum test results for differential gene expression comparing myocarditis versus healthy controls in clusters 0, 3, 1, 4, respectively, are shown in (**F**). Genes highlighted in red, or blue have adjusted p-values < 0.05.

To understand the functional role of myeloid cells in viral myocarditis, we compared gene expression profiles between groups, leading us to note 354 upregulated and 389 downregulated genes overall (**Table S3**). Among various subtypes of myeloid cells (**Fig 2D**), differences in gene expression patterns were noted in monocytes representing clusters 0, 3, and 4 and in macrophages in cluster 1 (**Fig 2F**). We then sought to understand the significance of differentially expressed genes in each cluster by focusing on genes whose log-fold change (logFC) expressions were ≥0.25 with adjusted p value <0.05, as well as a greater than 10% difference in percent of cells expressing the gene (Δ Percent of Cells) in myocarditic mice as compared to controls. Based on these criteria, in monocyte cluster 0 of myocarditis, we saw the upregulation of M2 macrophage marker genes *Ccl24*, *Arg1*, *Gatm*, and *Chil3,* which play roles in anti-inflammatory functions and fibrosis/tissue repair^21, 22^. Similarly, expression of *Tgfbi* and *S100a4,* which mediate tissue repair and survival of cardiomyocytes^23, 24^, may signify inhibitory functions of this cluster. *Ccl2* and *Ccl9,* markers of the M1 phenotype implicated in myocardial infarction ^25, 26^, were also found to be upregulated, thus suggesting a mixed phenotype for monocyte cluster 0 in myocarditis. Likewise, monocytes of cluster 3 in myocarditis had upregulation of *Ccl24* and *Ccl2,* in addition to *Lgals1* and *Wfdc17* which counter inflammation^27, 28^, suggesting that blood-derived monocytes arriving at inflamed hearts may take part in the reparative process. Similar analysis of the monocytes in cluster 4 revealed upregulation of mainly *Fcgr2b*, an inhibitory Fc receptor; *Ms4a6d*, a suppressor of IL-1b via NLRP3 activation; and *Gatm*, an activator of arginine metabolism, indicating their anti-inflammatory roles. Finally, transcripts in the macrophages of cluster 1 of myocarditis indicated upregulation of mainly *Ccl24, Chil3* (M2 markers), and those involved in the modulation of macrophage functions (*Fcgr2b*) and inflammation and/or cardiac remodeling (*Mt1, Wfdc17* and *Socs3*)^27, 29, 30^ (**Fig 2F**).

We next performed gene ontology (GO) analysis using the upregulated genes of myeloid cell populations in myocarditic mice. The analysis revealed prominence of metabolic pathways (oxidative phosphorylation, ATP, and TCA cycle), in addition to inflammatory responses, leukocyte migration and activation, hypoxia, and antigen-presentation functions (**Fig S2B**). Similar analysis corresponding to three clusters in myocarditis (0, 1 and 3) showed inflammatory and wound healing pathways to be highly prominent in the monocytes of cluster 0, whereas pathways related to cardiac muscle contraction, neutrophil degranulation and hypoxia were prominent in cluster 3 (**Fig S2C**). In contrast, macrophages of cluster 1 in myocarditis were related to mainly signaling molecules and negative regulators of inflammation and metabolic pathways (**Fig S2C**). Overall, scRNAseq analysis revealed M2 cells to be the major myeloid cells, with *Ccl24* being the major transcript to be upregulated in 3 out 4 clusters in myocarditis (0, 1, and 3) (**Fig 2F**), implying that *Ccl24* may be critical for M2 functions in myocarditic mice.

### Th17 cells form a dominant fraction of T cells in myocarditic mice

In dissecting the complexity of T cells, we identified nine subclusters (**Fig 3A**) by utilizing various phenotypic markers and unique transcriptional signatures (**Fig 3B, and Fig S3A**). Cells in each cluster varied, in that myocarditic mice had significantly higher proportions of T cells in clusters 1 (CD4^+^Th17), 2 (CD8^+^CTL), and 4 (CD4^+^Tregs), followed by 7 (CD4^+^*Tcf4*^+^), 6 (CD4^+^*Ki-67*^+^), and 5 (CD4^+^CD8^+^*Ccr1*^+^), whereas proportions of CD4^+^naïve T cells were higher in healthy mice, as expected (**Fig 3C**). However, no apparent differences in cell cycle status were noted between the two groups, since cells in most clusters in both healthy, and myocarditic mice were in the growing (G1) phase with minor variations between groups (**Fig S3B**). While predominance of T helper (Th) 17 cells and cytotoxic T lymphocytes (CTLs) indicate their effector functionalities, detection of Treg cells was not expected, and their presence indicates that Treg cells have a major role in the post-infectious phase of myocarditis. The significance of the presence of *Tcf4*^+^CD4 T cells is not clear, but *Tcf4* can promote NF-kB activation ^31^. Detection of CD4^+^*Ki-67*^+^ cells indicate their proliferating status, and the presence of CD4^+^CD8^+^ cells expressing *Ccr1* may suggest a possible role for them in disease mediation.

**Fig 3:**
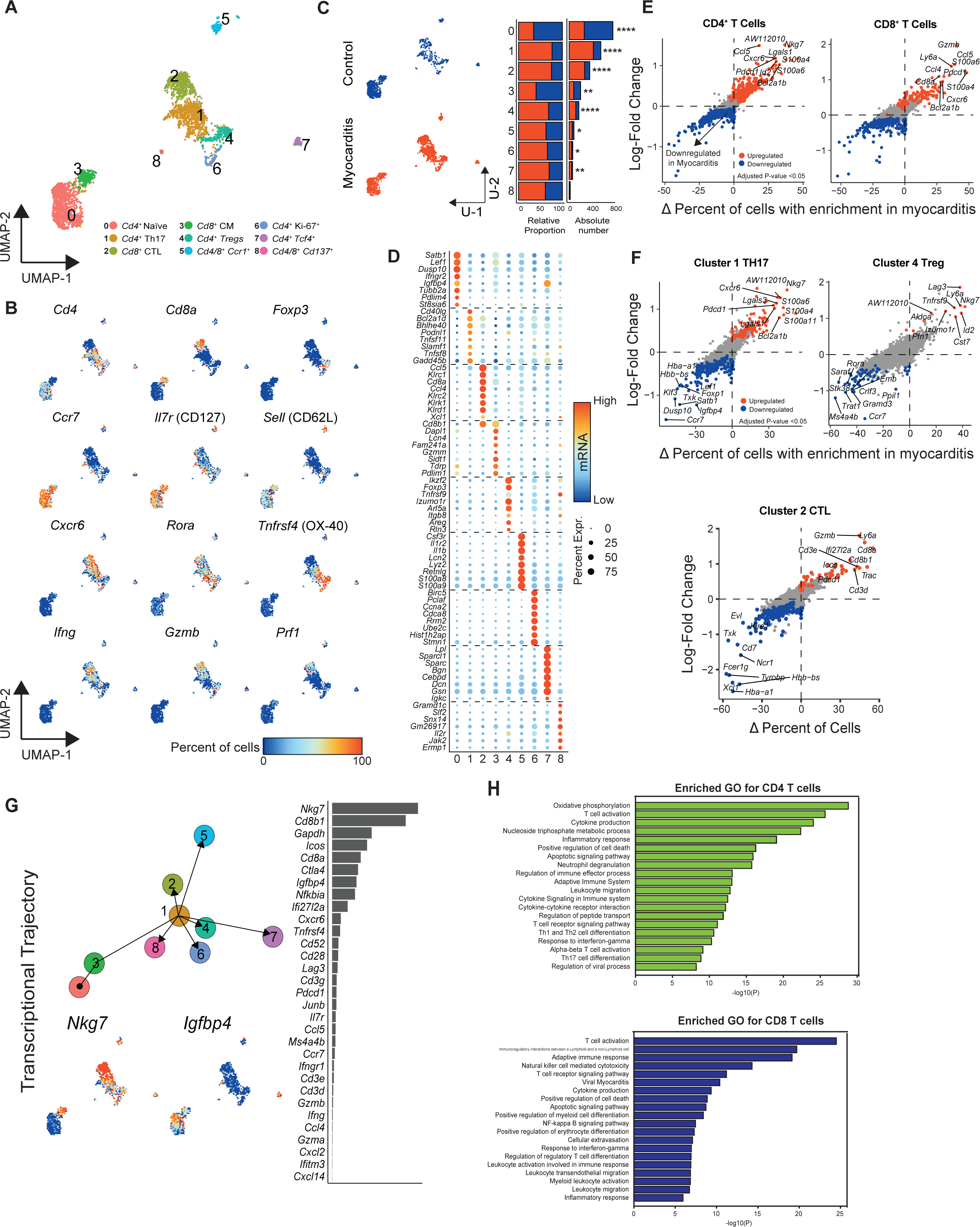
Analysis of T cell clusters reveals Th17 cells, CTLs, and Tregs to be the dominant fraction in myocarditis. Nine clusters were identified corresponding to healthy and myocarditic mice (**A),** using select markers for the indicated T cell types (**B)**, and their proportions relative to the total number of cells per condition are shown. Red indicates myocarditis, and blue indicates control. (**C**). Based on the mRNA expression pattern and log-fold change for the top eight transcripts in each cluster, a heatmap was then created, in which dot size equates to the percent of cells expressing the gene, while color corresponds to the expression level (**D**). Percentage difference (Δ Percent of Cells) and log-fold change based on the Wilcoxon rank sum test results for differential gene expression comparing myocarditis versus healthy controls in CD4^+^ and CD8^+^ T cells (**E**), and Th17, Treg, and CTLs (**F**), respectively, are shown. Genes highlighted in red, or blue have adjusted p-values < 0.05. Cell trajectory using the slingshot method ^19^ with Clusters 0 and 3 as the origin is shown; the top two genes with divergent expression at the root (*Igfbp4*) and at the terminal branches of the trajectory (*Nkg7*) are indicated in the UMAP (**G**). Ranked bar graph indicates the top immune-related genes in the order of their abundance in expression. GO analysis with enriched GO terms for various pathways (**H**) is shown for CD4^+^ (top panel) and CD8^+^ T cells (bottom panel).

We next identified the top eight differentially expressed genes in each T cell cluster across both healthy and myocarditic mice (**Fig 3D**). To understand their significance in viral myocarditis, we performed DGE on CD4 and CD8 T cells (**Fig 3E, Fig S3C, Table S3**), which revealed detection of upregulated genes that mediate various functions. For example, in myocarditic mice, we noted the upregulation in CD4 T cells of *Ccl5,* which is a target for NF-kB activation ^32^, whereas *Cxcr6* mediates recruitment of Th17 cells and CD8 T cells ^33, 34^. Strikingly, detection of *Nkg7,* which is implicated in cytotoxic functions ^35^, may mean that a proportion of CD4 T cells infiltrating hearts in viral myocarditis may have a cytotoxic function that has not been investigated thus far. Likewise, enhanced expression of *Pdcd1* and *Id2* may indicate the existence of potential checkpoints for T cell functions, since they are implicated in T cell exhaustion ^36^ and plasticity of Treg cells ^37^, respectively. Interestingly, DGE in CD8 T cells of myocarditic mice revealed increased expression of *Ccl5, S100a6, S100a4, Pdcd1, Cxcr6,* and *Bcl2a1b* similar to CD4 T cells, implying their common functionalities in both CD4 and CD8 T cell subsets (**Fig 3E**). While expression of *Gzmb* validates the identity of CD8 T cells, *Ly6a*, a regulator of memory T cell development, is also expressed in CD4^−^CD8^−^Treg cells ^38^ and may have a role in viral myocarditis.

Since T cell infiltrates in myocarditic mice had a predominance of Th17 cells, Treg cells, and CTLs (**Fig 3C**), we sought to analyze gene expression profiles in these subsets, expecting to identify novel genes of interest that may be involved in the viral pathogenesis. Th17 cells in myocarditic hearts when compared to those in controls (**Fig 3F, top left panel**) showed elevated expression of genes that were also upregulated in the whole CD4 T cell DGE analysis (**Fig 3E**). While expression of *Cxcr6* was expected in Th17 cells, upregulation of *Nkg7* and *S100a4*, which respectively mediate cytotoxic function^35^ and tissue repair^23^, was not expected (**Fig 3F**). The Treg cells in cluster 4 of myocarditic mice (**Fig 3F, top right panel**) had upregulation of their known markers *Lag3, Ly6a, Tnfrsf9, Izumo1r, Id2, Cst7*, and fructose-bisphosphate aldolase, along with increased expression of *Nkg7* and *Pfn1,* which mediate cytotoxicity^35, 39^, indicating that the cytotoxic Treg cells may be critical to maintaining homeostasis in the local inflamed heart milieu. Similarly, CTLs of cluster 2 in myocarditic mice (**Fig 3F, bottom panel**) indicated expected upregulation of *Gzmb*, CD8 coreceptors, and Cd3 complex proteins; elevations of *Icos* and *Pdcd1* may indicate activation status of CTLs ^36, 40^. We next built cell trajectories using Slingshot (**Fig 3G**). All T cell clusters had a common root-point from CD4^+^naïve cells (cluster 0) together with CD8^+^ CM cells (cluster 3), branching into CD4^+^Th17 (cluster 1), which gave rise to terminal branches of other T cell populations (**Fig 3G**). We noted two genes with divergent expressions at the root (*Igfbp4*) and at the terminal branches (*Nkg7*) (**Fig 3G**). *Igfbp4* can mediate both proliferative and inhibitory functions, whereas *Nkg7* is implicated in cytotoxic functions ^35^. The finding that upregulated expression of *Nkg7* was evident in the whole CD4^+^ T cell subset (**Fig 3E**), Th17 cells, and Treg cells in myocarditis (Fig 3F, top panel), suggested the possibility that *Nkg7* may have a role in the effector functions of these cell types in viral myocarditis. Furthermore, gene set enrichment analysis (GSEA) indicated that the genes in the whole CD4 T cell subset were mostly involved in immune metabolism, T cell activation, Th1, Th2 and Th17 differentiation, apoptosis, and cytokine production, whereas T cell activation, cytotoxicity, apoptosis, and NF-kappa B signaling and inflammatory response-related pathways were apparent in CD8 T cells of myocarditic mice (**Fig 3H**). These observations were also recapitulated individually in Th17 cells and CTLs of myocarditic mice (**Fig S3D**). Together, the data suggest prominent roles for Th17 cells, Tregs, and CTLs in the immune pathogenesis of viral myocarditis, with a possibility that the effector functions are mediated mainly by cytotoxic functions regardless of T cell subsets.

### Fibroblasts in myocarditic mice contained various subtypes with unique transcriptome signatures that can influence functionalities of immune cells

We analyzed the fibroblast population based on the known markers (**Fig 1E**) and re-clustered it into 12 distinct populations as indicated in the UMAP (**Fig 4A**). We analyzed the relative proportion of cells in each cluster and noted the cells in clusters 1, 5, 6, and especially 8, were present in greater numbers in myocarditic mice than in healthy mice (**Fig 4B**). Additionally, we noted that fibroblasts in myocarditic mice tended to be in the S phase compared to healthy mice, and such a trend was more evident in cluster 8 than others (**Fig S4A**). We next analyzed gene expression profiles of the top eight differentially expressed genes in all 12 fibroblast populations, revealing a few noteworthy findings. Expression of enriched genes, although unique in all clusters, showed overlapping patterns in clusters 5, 6, and 8 (**Fig 4C**). We noted three genes, *Postn* (periostin), *Ltbp2* (latent TGF-β binding protein 2), and *Thbs4* of interest in cluster 5. While *Postn,* an ECM protein, is critical for tissue development and regeneration and plays a role in wound healing and ventricular remodeling in myocardial events ^41^, *Ltbp2* and *Thbs4* have roles in myocardial fibrosis in DCM ^42^ or the fibrotic process^43^. Similarly, three transcripts, *Apoe*, *Tgfbi,* and *Mfap4,* were found interesting in cluster 6. They mediate regulation of T cell and macrophage functions, including inflammation and oxidative stress ^44^, cardiac fibrosis in concert with *Postn* ^24^, and ventricular remodeling and cardiac function, respectively ^45^. As to cluster 8, we noted that a few genes were uniquely expressed with varied functions. These include *Apoe, Tgfbi*; *Wif1*, which modulates cardiomyocyte differentiation ^46^ necessary for cardiac remodeling in myocardial infarction and also promotes DCM ^47^; *Clu*, myocardial injury marker ^48^; and *Npy,* which has a role in cardiac remodeling and heart failure ^49^. These data indicated that clusters 5, 6, and 8 were mainly involved in the cardiac remodeling process in myocarditis. Although there also were more fibroblasts in clusters 1 and 3 in myocarditic mice, their transcriptional profiles were unique to each (**Fig 3C**). The genes expressed in cluster 1 include *Mt1* and *Mt2,* which can modulate inflammation and support cardiac remodeling as demonstrated in ischemic cardiomyopathy ^30^; *Thbs1*, a promoter for *Tgfbi* activation in fibrosis ^50^; *Cxcl1*, which facilitates recruitment of neutrophils and non-hematopoietic cells to the site of injury and regulates immune and inflammatory responses ^25^; and *IL-6*, inducer of MMP1 that can mediate tissue remodeling ^51^. Regarding cluster 3, expressions of *C3* (activator of complement system), and *Apod* (putative marker of non-proliferating and senescent fibroblasts) ^52^ were elevated. These observations suggest that upregulated genes in the corresponding clusters have a role mainly in cardiac remodeling events and immune activation. To further determine the specific roles of potential transcripts that might contribute to CVB3 pathogenesis, we performed DGE in fibroblast clusters 1, 5, 6, and 8 of myocarditis and control (**Fig 4D**). These analyses revealed increased expression of complement proteins *C3* and *C4b*; *Serpina3n,* an accelerator of wound healing ^53^; *Tmem176b*, which controls DC maturation and also has a role in fibrosis ^54^; *Tmem176a* a negative regulator of DCs ^55^; and the anti-viral protein *Ifitm* ^56^ in all clusters of myocarditic mice (**Fig 4D**). Uniquely however, *Sbno2*, a regulator of proinflammatory cascade ^57^ and *Ly6a,* also called *Sca-1*, which has a known function in heart failure ^58^, were increased in cluster 1 of myocarditic mice, as opposed to *Gsta3,* an inhibitor of TGF-β-induced epithelial and mesenchymal transition and fibronectin expression ^59^, and *Socs3* ^29^ in cluster 5; *Apod* ^52^ in cluster 6; and *Apoe* in cluster 8 of myocarditic mice (**Fig 4D**).

**Fig 4:**
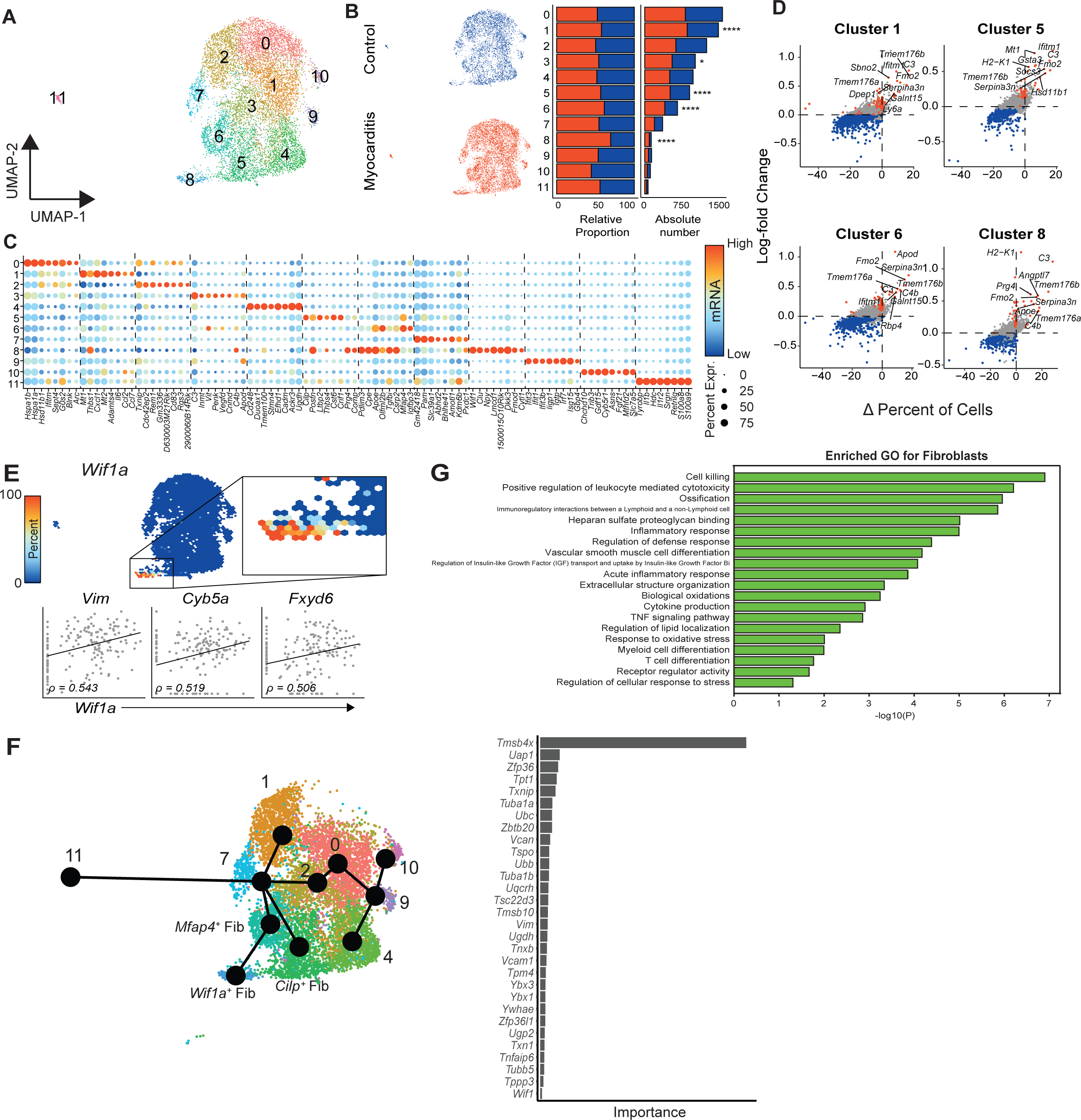
Analysis of fibroblasts reveals new cardiac fibroblasts involved in cardiac remodeling during viral myocarditis. (**A**) indicates the UMAP of isolated fibroblasts across groups; their relative distributions in healthy and myocarditic mice are shown in UMAPs and cluster-wise in bar plots, with red indicating myocarditis, and blue indicating control (**B**). Panel (**C)** represents the heatmap of the top eight markers shown with log-fold change, where dot size and color represent percent of cells expressing the gene and expression levels, respectively. Panel (**D**) indicates differential expression of the top 10 genes in myocarditic mice as compared to healthy mice with respect to clusters 1, 5, 6, and 8. Percentage difference (Δ Percent of Cells) and log-fold change are based on the Wilcoxon rank sum test results. The panel (**E**, top panel) shows localization of *Wif1* expression in fibroblast cluster 8, and its correlation with other markers *Vim, Cyb5a,* and *Fxyd6* (**E**, bottom panel). Slingshot analysis along the UMAP (**F**) shows the root and terminal branches of the cell fates of fibroblast populations. Branches terminating in *Wif1a*^+^ fib and *Cilp*^+^ fib cell clusters promote cardiac remodeling. GO analysis with enriched GO terms for various pathways (**G**) is shown for the whole fibroblast population.

Since there were more fibroblasts in cluster 8, which also expressed injury marker *Wif1a*, we sought to correlate its expression with other genes (**Fig 4E, top panel**). This analysis revealed a positive correlation with *Vim*. However, unexpectedly, two other genes (*Cyb5a* and *Fxyd6*) also showed positive correlation with *Wif1a* expression (**Fig 3E, bottom panel**); whether their expression can also be used as injury markers remains to be investigated. We next explored the origins of cardiac remodeling-associated fibroblasts by analyzing transcriptional activation using Slingshot. The analysis suggested that *Cilp*^+^ cluster 5 had its origins in cluster 7, whereas *Wif1a*^+^ cluster 8 arose from cluster 6, which in turn had its origins in cluster 7 (**Fig 4F, left panel**), with *Tmsb4x,* having roles in repair of human heart muscle, being the important transcriptional activator for this cell trajectory (**Fig 4F, right panel**). Finally, by performing GO analysis on all populations, we noted the upregulation of functions associated with inflammatory responses and other immune signaling pathways (**Fig 4G**), including cytotoxicity, ossification, ECM protein synthesis, and immune/inflammatory regulatory networks, among others (**Fig 4G, and Fig S4C**), suggesting that fibroblasts can perform diverse functions during CVB3 viral myocarditis.

### NK cells, ILC2, and ILC3 cells formed a major component of ILCs, but their numbers were low in myocarditic mice

We analyzed ILCs pooled from healthy and myocarditic mice, and the use of canonical markers allowed us to dissect ILCs into five distinct populations (**Fig 5A**). As shown in Fig 5B, three markers of NK cells (*Nkg7*, *Klrd1,* and *Gzma)* were consistently expressed in clusters 0, 1, and 3. Conversely, ILC markers (*Gata3, Ltb4r1, Csf2* and *Il17a*) were expressed in clusters 2 and 4. By evaluating the relative proportion of cells present in each cluster, we observed that the Gzma^+^ NK cells, Tcf7^+^ NK cells, and Gata3^+^ ILCs were significantly reduced in myocarditic mice as compared to healthy controls (**Fig 5C**). Next, we evaluated the DGE in NK cell clusters and identified the top 15 differentially expressed genes between groups (**Table S3**). **Fig 5E and Fig S5A** shows upregulation of genes involved in NK cell development (*Ltb* and *Thy1*) ^60^ and NK cell function (*Gzmc*). Genes implicated in cell migration, invasion, and fibrosis (*S100a4* and *S100a6)*^23^, and anti-viral function (*Ifi27l2a*)^56^ were also upregulated in myocarditis. While upregulation of *Pdcd1* (PD-1) may represent activation status, *Cxcr6* expression may indicate that NK cells or ILCs infiltrating the hearts can have effector functions as noted with HIV and VSV infections ^61^. Another transcript, *Tmem176b,* may be a novel candidate with a role in cardiac fibrosis since upregulation of this gene has been noted in pulmonary fibrosis ^54^. By contrasting these patterns in individual clusters, it was clear that only *Gzma*^+^ NK cells in cluster 0 had a similar profile in both myocarditic and control mice, and to a lesser extent the *Tcf7*^+^ NK cells of cluster 1 (**Fig S5B**). Finally, the GO analysis found that all NK cell clusters had a role in the activation and regulation of T cells and neutrophils and anti-viral responses in myocarditis (**Fig 5F**), with similar functions being noted in *Gzma*^+^ NK cells (cluster 0) in addition to the apoptotic signaling pathways (**Fig S5C**). Together, the data suggest a role for infiltrating NK cells in myocarditic mice in post-infectious myocarditis, but with no or minimal contribution from ILCs.

**Fig 5:**
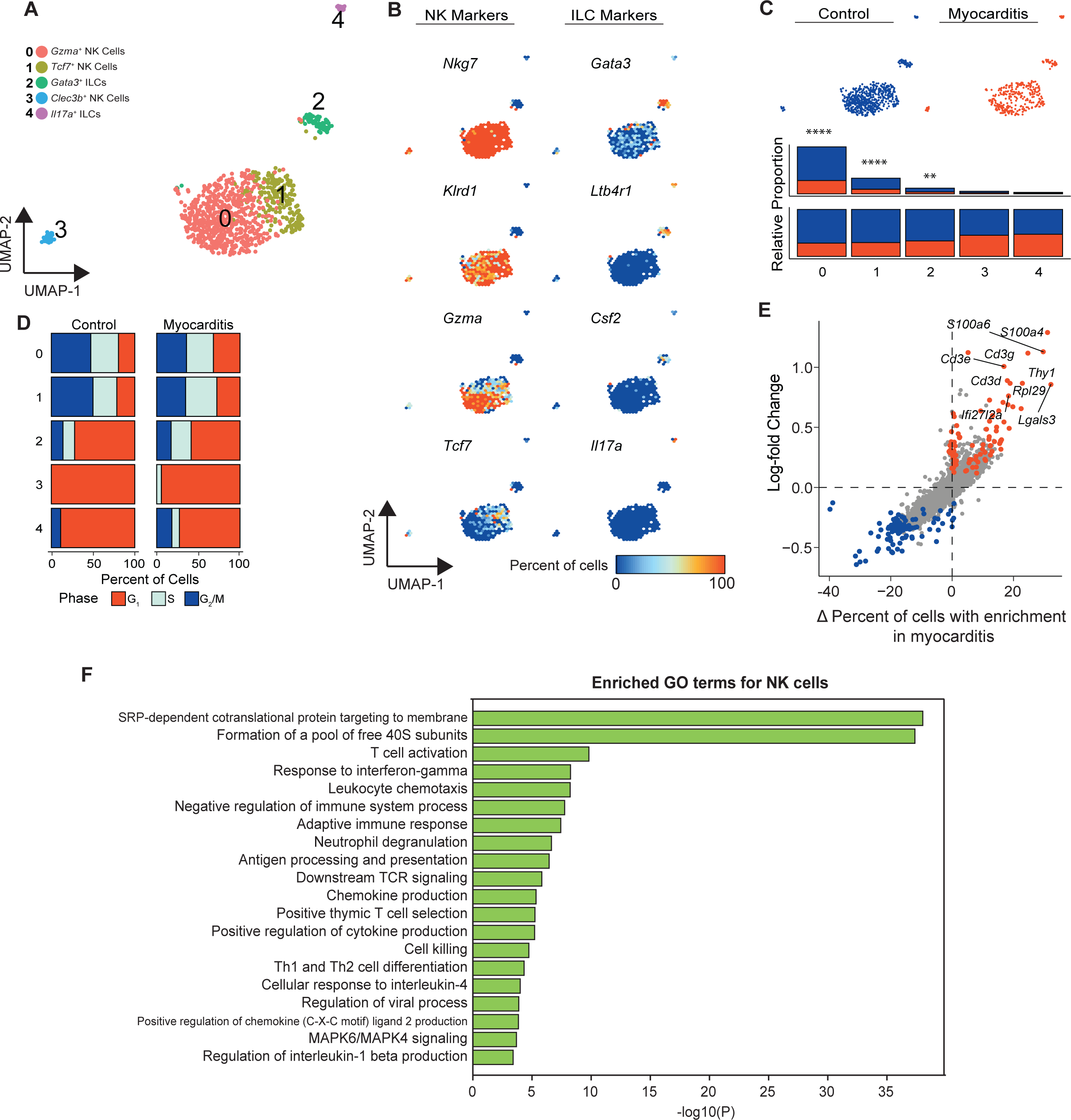
Analysis of ILCs in heart infiltrates of myocarditic mice. (**A**) indicates the UMAP of ILCs across groups; the canonical markers used to identify the five clusters of ILCs are shown in (**B**). After their distributions corresponding to healthy and myocarditic mice were identified (**C, top panel),** the proportion of cells in each cluster relative to the total number of cells per condition was determined, with red indicating myocarditis, and blue indicating control (**C, bottom panel**). The panel (**D)** indicates cell cycle assignments across all ILC and NK cell clusters in healthy controls and myocarditic mice, Χ^2^ test p-value < 2e-16; in panel (**E)**, the percentage of differentially expressed transcripts (Δ Percent of Cells and log-fold change) in NK cells in myocarditic mice is shown. Genes highlighted in red, or blue have adjusted p-values < 0.05. Panel (**F**) indicates enriched GO terms with respect to pathways upregulated in NK cells of myocarditic mice.

### Neutrophils infiltrating the myocarditic hearts predominantly modulate inflammatory responses

By utilizing the known markers, we subclustered the neutrophils obtained from myocarditic and healthy mice into six populations (**Fig 6A, Fig S6A**). All clusters had similar proportions of neutrophils in both groups, barring *Ly6g*^+^ cluster 2 and *Ccl5*^+^ cluster 6. Whereas the cells in myocarditic hearts in cluster 2 were lower than in the healthy control, an opposite trend was noted in cluster 6 (**Fig 6B**). Evaluation of the top eight differentially expressed genes in each cluster revealed *Camp*, an antimicrobial peptide; and *Ly6g*, a marker of myeloid cells/granulocytes, as highly expressed genes in *Ly6g*^+^ cluster 2, whereas *Ccl5*, a chemokine expressed by non-neutrophils^25^, was identified in cluster 6. Similarly, increased expressions of *Lyz2*, a chemotactic for neutrophils; and *Ccl6*, a chemokine expressed by neutrophil and macrophage lineages^25^ were seen in *Ccl6*^+^ cluster 0. In addition, *Fgl2*, a regulator of immune and inflammatory responses^62^; and *Itgax* or *CD11c*, a marker of DCs that triggers respiratory burst in neutrophils, were found in *Gm2a*^+^ cluster 1 (**Fig 6C**). Interestingly, cluster 3 neutrophils contained genes involved in the regulation of inflammation via inhibition of NF-kB, including *Nfkbia, Nfkbie*, *Tnfaip3,* and *Tnf*. Likewise, *Ifit1*^+^ cluster 4 had genes related exclusively to anti-viral activity (*Ifi1, Ifit3b* and *Oasl1*) ^56^, whereas those in *Mt-High* cluster 5 were all mitochondrial genes (*Mt-nd1/2/3* and *Mt-atp6*) that have been implicated in mitochondrial cardiomyopathy ^63^. The data thus suggest a heterogeneity in neutrophil populations that mediate varied functions in viral myocarditis.

**Fig 6:**
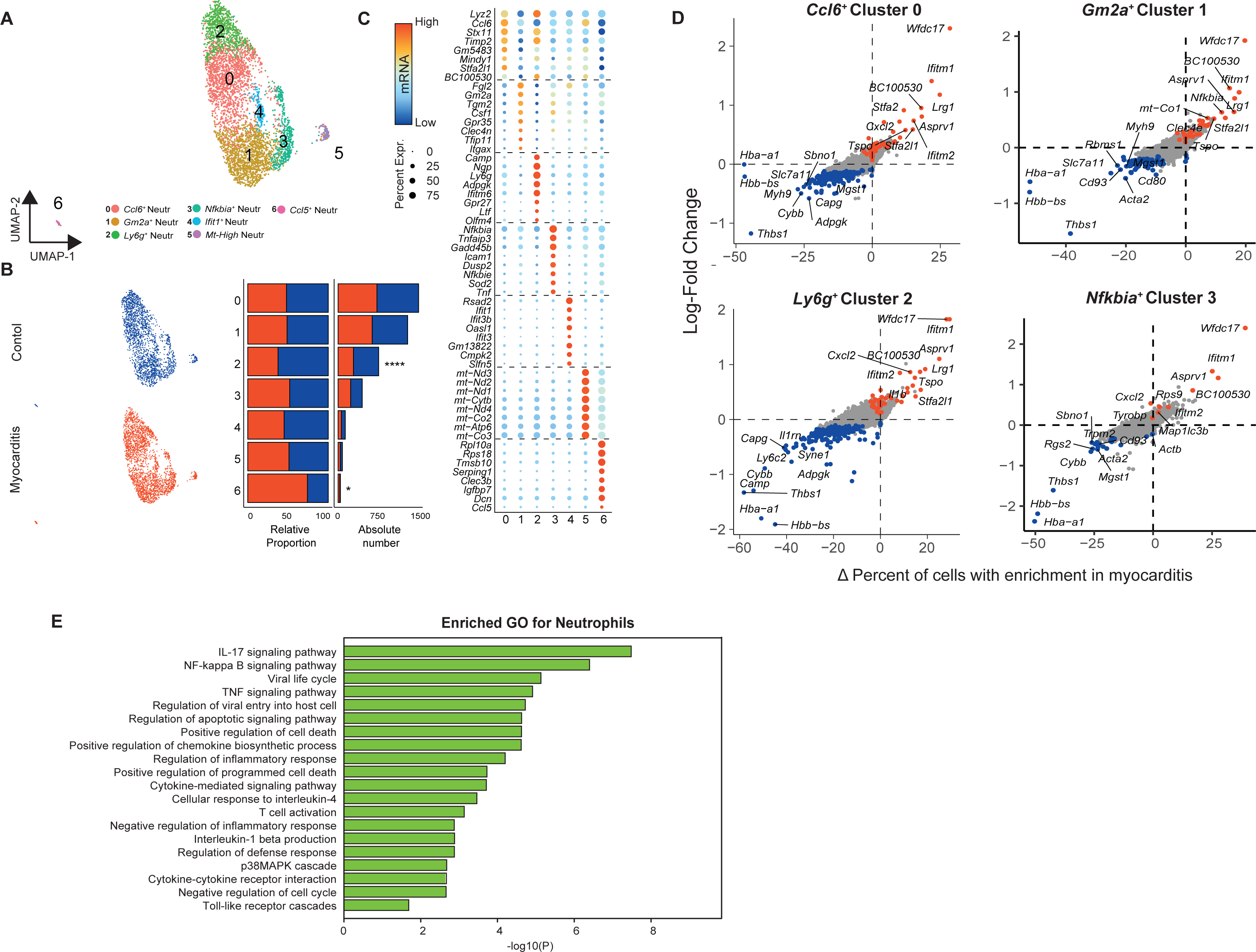
Neutrophils mainly with pro-inflammatory functions were detected in myocarditic mice. **A**. UMAP of neutrophils representing seven clusters across groups. **B.** UMAPs and bar plots demonstrating the relative distribution of neutrophils in healthy and myocarditic mice by cluster, with red indicating myocarditis, and blue indicating control. **C**. Heatmap of the top eight markers by log-fold change. Dot size equates to the percent of cells in the cluster expressing the gene, while color corresponds to the expression level. **D**. Percentage differences (Δ Percent of Cells) vs. log-fold change of the differentially expressed genes in myocarditic mice relative to the healthy group are indicated for clusters 0,1, 2, and 3. **E**. Enriched GO terms with respect to pathways upregulated in neutrophils of myocarditic mice.

Through DGE analysis in the myocarditic mice, we also noted a pattern that had similarities in several clusters. For example, prominent upregulated genes in *Ccl6^+^*cluster 0 of myocarditis included *Wfdc17*, negative regulator of inflammation; anti-viral proteins *Ifitm1, Ifitm2*, and *Lrg1*, a novel neutrophil granule protein and modulator of myelopoiesis that also has a role in wound healing; and *Tspo*, a mitochondrial membrane protein overexpressed in inflammatory processes (**Fig 6D**). Similar trends were noted in *Ly6g*^+^cluster 2 in myocarditis, except that *Il1b* was upregulated, which is known to promote cardiac fibrosis and remodeling in myocardial infarction^64^ (**Fig 6D**). Likewise, *Clec4*, an interacting partner of *FcRIγ*, was increased in cluster 1 of myocarditic mice (**Fig 6D**). While gene expression profiles in *Nfkbia*^+^ cluster 3 were similar to those in clusters 0 and 2, no differentially expressed transcripts were noted in clusters 4 to 6 (data not shown). We next performed GO analysis, leading us to note that the pathways involved in the regulation of various immune and inflammatory responses (*Il17*, NF-kB, TNF-signaling), cell death, cytokine signaling (*Il1β*, *Il4* signaling), as well as modulation of viral life cycle, were upregulated in myocarditis (**Fig 6E**). Several of these pathways also overlapped across individual neutrophil clusters, especially in cluster 1 of myocarditic mice (**Fig S6B**); Cluster 0 predominantly showed pathways of interferon functions, and cluster 2 showed neutrophil chemotaxis and inflammatory response. In sum, the data showed that neutrophils participate in the pathogenic inflammatory and cardiac fibrosis process, with *Il1β* being the major driver of this process.

### Other cell types detected in myocarditic mice included B cells and, to a lesser extent, ECs, basophils, and SMCs

As presented in **Table S2**, scRNAseq analysis revealed the identification of cells other than myeloid cells, T cells, fibroblasts, or NK cells/ILCs in varied proportions. These included B cells, basophils, ECs, erythroid cells, and SMCs. Of these, we did not investigate the transcriptomes of basophils, ECs, and SMCs because their numbers were low (**Table S2**). We also did not consider erythroid cells for downstream analysis since they were not our major focus. However, by using known markers of B cells, we analyzed the B cell population, leading us to identify five distinct cell populations (**Fig S7A**). After evaluating the relative proportion of cells in each cluster, we noted that ‒ except for cluster 4 ‒ cell proportions were reduced in all clusters, although not significantly in myocarditic hearts (**Fig S7B**). Nonetheless, by analyzing the differentially expressed transcripts, we found no significant differences between groups (**Fig S7C**), suggesting that the low proportion of B cells that accumulated in the myocarditic mice appeared not to play a major role in the immune pathogenesis of CVB3 infection.

### Ligand-receptor analysis revealed diverse intercellular communications during myocarditis

To determine the intercellular communication networks between various cell types, we used CellChat ^65^, which utilizes the known structural composition of 2,021 validated ligand-receptor interactions and membrane-bound co-receptors deposited in CellChatDB ^65^. While communications were evident between all cell types detected in both myocarditic (**Fig 7A**) and healthy (**Fig S8A**) mice, the number of ligand-receptor pairs involved in these interactions differed between groups as indicated by dense intercellular communication networks. Notable among these, especially in myocarditis, were the following (ligand-receptor): B cells-CD8 T cells; basophils-CD8 T cells and myeloid cells; ECs-CD8 T cells; CD4 T cells-CD8 T cells; CD8 T cells-neutrophils, myeloid cells, and NK cells; fibroblasts-SMCs and myeloid cells; and ILCs-CD8 T cells (**Fig 7B and Fig S9**). CellChat also allowed us to evaluate the differential number of interactions and their strength in the myocarditic cardiac cellulome, as compared to healthy cells, based on the upregulated ligand-receptor pairs. We made a few observations: 1) The interaction strength and number of signals sent from fibroblasts to CD4 and CD8 T cells and SMCs were enriched in myocarditis (**Fig 7C and Fig S8B**). 2) Myeloid cells appeared to have stronger and more interactions with CD4 T cells and SMCs. 3) The number of SMC interactions between most of the cell types and their relative strength were enriched. However, the analyses revealed no indication of autocrine networks for fibroblasts and myeloid cells (**Fig 7A, 7C, and Fig S8B**). GO enrichment analysis of upregulated interactions between cell types revealed that the ligand-receptor pairs involved in fibroblasts, CD4 and CD8 T cells, were associated with positive regulation of proliferation of epithelial cells and SMCs, adhesion and migration of cells, and inflammatory cytokine responses (*IL-1β, IFN-γ and TNF-α*) (**Fig 7D, 7E and Table S4**). In combination with fibroblasts, ligand-receptor pairs involved in myeloid cells were also associated with cell adhesion and anti-inflammatory and wounding and regeneration processes. However, fibrosis-associated processes were mainly restricted to the receptor-ligand pairs in the fibroblasts, suggesting a prominent role for them in the development of cardiac fibrosis in viral myocarditis. We next identified 66 signaling pathways associated with the ligand-receptor pairs we had identified, and we noted these to be enriched in either myocarditic (EGF, CD52, CXCL, CCL, MHC-I, LCK, ncWNT, AGRN, OSM, NOTCH, VTN, PDGF, GRN and CD45) or healthy mice (MK, THY1, PTN, CD80, CD23, CDH1, NECTIN, CLEC, EPHB, VISFATIN, IL2, MPZ, among others), whereas a few others were enriched equally in both (**Fig 7F**). Notably, most of the pathways enriched in myocarditic mice, such as EGF, LCK, ncWNT, OSM, NOTCH, and PDGF, have been implicated in CVB3 pathogenesis or atherosclerosis/cardiac fibrosis. For example, in our data we observed that the EGF signaling involved in pathogenic plaques and remodeling of blood vessels^66^ was upregulated in myocarditis, with signals being sent from the ILCs and SMCs to the fibroblasts (**Fig S10**). LCK signaling of the Src family p56^lck^, which is essential for CVB3 replication and pathogenesis^67^, had signaling interactions from NK cells and CD4 T cells to CD8 T cells, with additional strong autocrine signaling on CD8 T cells (**Fig S10**). For the ncWNT pathway, which plays a role in cardiac remodeling^68^, signals were being transmitted from SMCs to ECs, with autocrine signaling on SMCs (**Fig S10**). OSM, which has been found to be upregulated in DCM patients^69^, included signals sent from basophils, myeloid cells, and neutrophils to SMCs (**Fig S10**). NOTCH signaling, which plays a role in the repair of the myocardium^70^, was secreted from SMCs in an autocrine fashion and to the ECs and neutrophils (**Fig S10**). The profibrotic signaling in PDGF was secreted by the ECs and SMCs to fibroblasts, in an autocrine network, as well. Finally, CD45 signaling was strongly secreted from almost all cells to the myeloid cells (**Fig S10**). Overall, the intercellular communication patterns in the cardiac cellulome and the signaling pathways being prominently upregulated in myocarditic mice may contribute to and play a critical role in the pathogenic progression of viral myocarditis.

**Fig 7:**
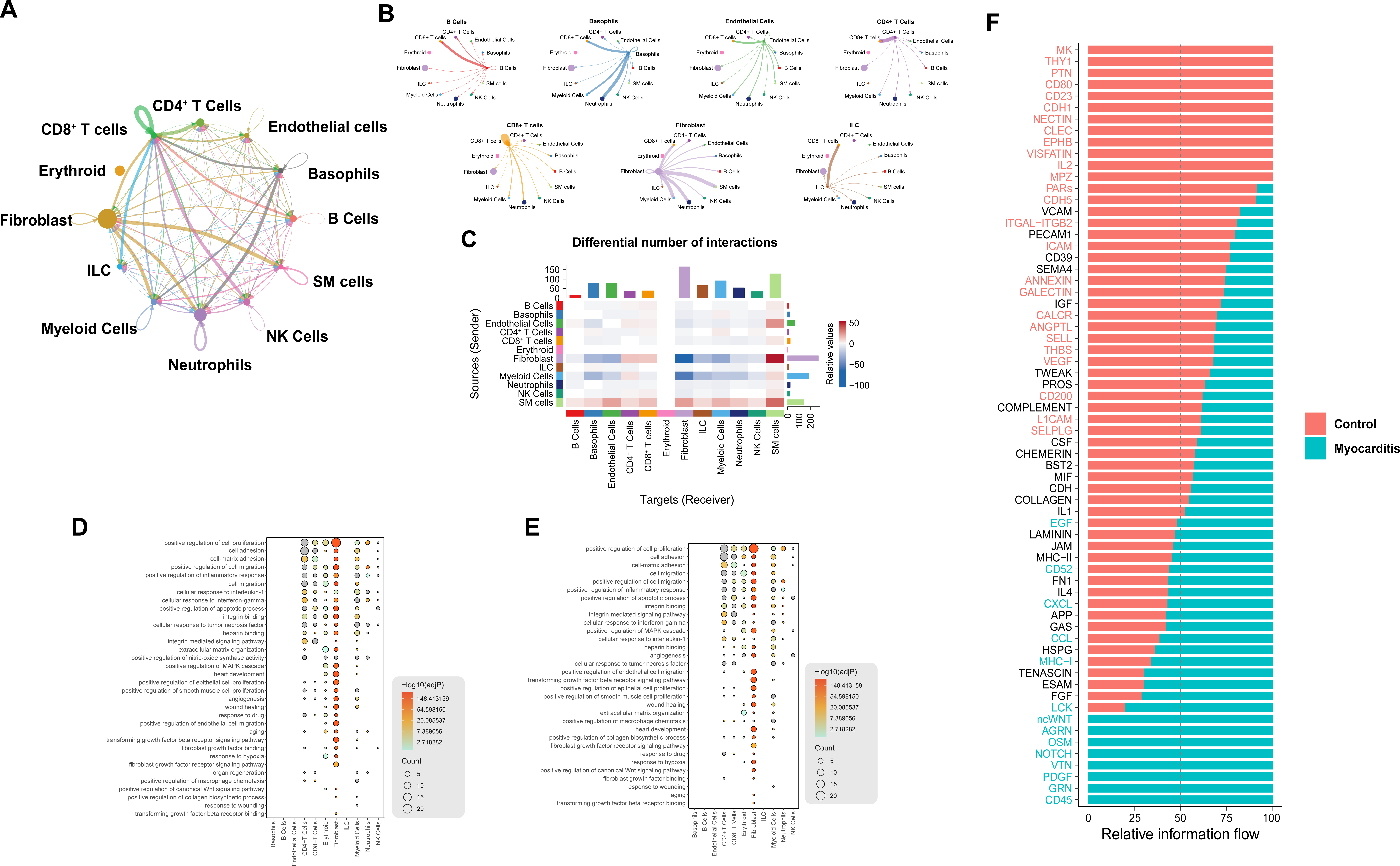
Intercellular communication between cardiac cell types in myocarditis. **A**. Circle plot showing the intercellular communication between major cardiac cell types, using CellChat R workflow. The lines originating from a cell type indicate ligands being broadcast, with these lines connecting to the cell type where the receptors are expressed. Thickness of the line is proportional to the number of unique ligand-receptor interactions, with loops representing autocrine circuits. **B**. A detailed view of ligand and cognate receptor interaction for major cell types. **C.** Heatmap of differential number of interactions between myocarditic and healthy mice in the cell-cell communication network. The top-colored bar plot indicates the sum of column values (incoming signaling), and the right bar plot indicates the sum of row values (outgoing signaling). Red indicates increased signaling in myocarditis, and blue indicates decreased signaling. **D.** GO terms enriched in a set of genes that encode ligands upregulated in myocarditis. GO terms are ordered by their frequency of significant enrichment in different cardiac cell populations. **E**. GO terms enriched in a set of genes that encode receptors upregulated in myocarditis. GO terms are ordered by their frequency of significant enrichment in different cardiac cell populations. Dot size indicates number of enriched genes, with colors indicating -log10 (adjP) value. **F.** All significant signaling pathways ranked based on their relative information flow within the inferred networks between healthy and myocarditic mice. The top signaling pathways colored red are more enriched in control mice, the ones colored black are equally enriched in control and myocarditic groups, and the blue colored pathways are more enriched in myocarditic mice.

### TFs enriched in myocarditis modulate cardiac remodeling functions of target genes

In order to identify the TFs that regulate the differential expression of genes in different cell clusters, we employed single-cell regulatory network inference and clustering (SCENIC)^71^. This analysis revealed upregulated expression of four TFs in both CD4 and CD8 T cells (*Elf1, Ets1*, *Irf7*, and *Stat1*) (**Fig 8A**). Functionally, *Elf1* is known to regulate anti-viral responses, especially of Type I IFNs ^72^, and *Ets1* controls expression of cytokines and chemokines ^73^, whereas *Stat1* is involved in the signaling cascades of both Type I and Type II IFN responses. We then analyzed co-expression patterns of these TFs with the upregulated genes in the CD4 and CD8 T cell subsets (**Fig 8B, Table S5**). Among all TFs, we noted increased interactions mainly between *Ets1* and target genes involved in cardiac ischemia, remodeling, and heart failure (*Ccr5, Ccl5, Cxcr3, Ccr2, Cxcr6,* and *S100a4*)^23, 25^. As shown in **Fig S11A**, the target genes of *Elf1* and *Ets1* can mediate TCR signaling and T cell activation among other functions (cell adhesion, chemotaxis, and inflammation). Similarly, *Stat1*-targeted genes facilitate antigen-presentation/cross-priming, and immunoproteasome functions in myocarditic mice. By extending these observations to myeloid cells, we noted upregulation of five TFs implicated in various functions (**Fig 8A**). These include *Irf5, Mafb*, *Maff*, *Mef2c,* and *Rara*. The putative co-expression patterns showed increased interactions of *Irf5* and *Mafb* with each other, and with target genes involved in fibrosis and remodeling (*Tgfbi, Fn1, Ccl24, Ccr2, Ccr5, Ccl2, S100a4*) and M2-specific phenotype (*Gatm, Arg1, Mrc1*) with *Irf5*, and *Mafb*-targeted genes mediating various processes, such as antigen presentation, leukocyte activation, response to *IFN-γ,* and cytokine production or inflammation (**Fig S11B**). We also extended these analyses to other cell types of interest, neutrophils and NK cells. We noted upregulation of one TF in neutrophils (*Bcl3*, a regulator of cell proliferation) and one in NK cells (*Egr1*, a regulator of growth, cell survival and cell death) in myocarditic mice, but no apparent differences were noted between groups in fibroblasts (**Fig S12**). Overall, since T cells and myeloid cells form a major component of cellular distribution in myocarditic mice and also upregulate multiple TFs that can control various immunological processes involving many of the upregulated target genes in the respective cell types, the data point to a major role for them in the immune pathogenesis of CVB3 infection.

**Fig 8:**
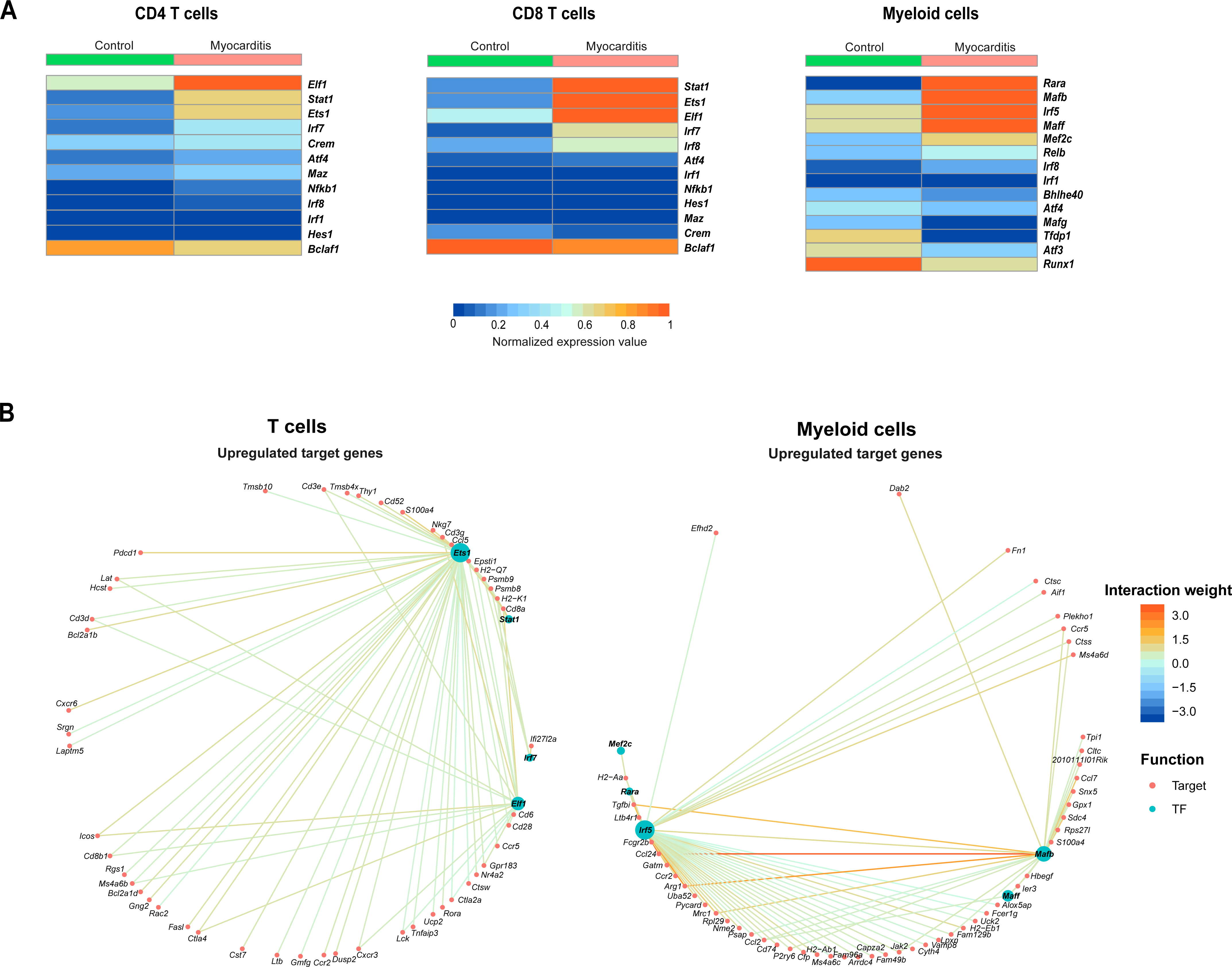
Analysis of myocarditis-specific transcription factors and their target genes. **A.** Heatmaps showing the transcription factors being enriched in either myocarditis or control hearts for CD4, CD8, and myeloid cells using SCENIC. **B.** Network plot showing the upregulated target genes for the indicated transcription factors in T cells and myeloid cells. Lines indicate the interaction between TFs and their target genes, and colors indicate the interaction weight range.

## Discussion

In this report, we have described the cellular complexity that occurs during the post-infectious phase of viral myocarditis induced with CVB3. Previous reports have delineated the cardiac landscape and intercellular communication in healthy mice ^74^ and have investigated the fibrosis and cardiac remodeling process in angiotensin II (*AngII*) mouse models of fibrosis ^75^. To our knowledge, this is the first report to comprehensively dissect the cardiac cellulome and the intercellular communication networks in viral myocarditis. By using whole heart cells, we were able to capture a majority of immune cells, including fibroblasts, and a fraction of the ECs and SMCs (**Fig 1F, G**), but not cardiomyocytes ‒ possibly because droplet-based sequencing techniques are unable to process large cells. Enrichment of the latter three cell types also require specialized protocols utilizing single nuclei isolation, which we did not use in our studies. After ascertaining the identities of various cell types, myeloid cells, T cells, and fibroblasts were found to be significantly increased in myocarditic mice as compared to healthy mice (**Fig 1G**).

In myeloid cells, we detected mainly monocytes and macrophages, and all subclusters of myeloid cells were branched from monocytes, with *Msrb1* being the top immune-related gene driving this differentiation of myeloid cells (**Fig 2D**). *Msrb1* is a selenoprotein promoting anti-inflammatory cytokine expression and has been found to be upregulated in mouse models of cardiac stress ^20^. The fact that this gene drives the differentiation of myeloid cells could indicate the anti-inflammatory role of these cells in the post-infectious myocarditis phase. In addition, by analyzing the transcriptomes, we noted upregulation of several genes that have roles in M2 macrophages and anti-inflammatory functions. Importantly, upregulation of *Ccl24* was consistently noted in both monocytes and macrophages (**Fig 2F**). *Ccl24*, also called eotaxin-2, was shown to promote pathogenic fibrosis in skin, lung, and liver models of fibrosis in mice ^22, 76^; it also was produced by F4/80^+^ macrophages in inflamed hearts, which may facilitate eosinophils in eosinophilic myocarditis ^77^. Although eosinophils are not commonly reported in viral myocarditis, *Ccl24* expression may be necessary to recruit monocytes or facilitate their conversion to M2 cells, which may be critical to repair damaged cardiac tissue or participate in cardiac fibrosis. Additionally, the upregulated genes of myeloid cells were found to have a prominent role in metabolic pathways, especially oxidative phosphorylation, ATP, and the TCA cycle (**Fig S2B, C**); the dependency of M2 cells on these pathways has been demonstrated in macrophages ^78^. Taken together, the myeloid cell populations noted in post-infectious myocarditis may primarily be involved in the reparative process in affected animals.

By investigating the T cell landscape, we noted few observations that offer new insights into myocarditis pathogenesis. T cells mainly consisted of Th17 cells, CTLs, and Tregs in myocarditic mice (**Fig 3C**). Detection of Th17 cells was not surprising, since IL-17 blockade can ameliorate the severity of CVB3 myocarditis ^79^, but their antigen specificity remains unknown. This is critical because IL-17-deficient mice develop acute myocardial inflammation in the setting of autoimmune myocarditis, but they chronically develop DCM. It may be that a proportion of these T cells are specific to cardiac antigens, as we have demonstrated with MHC class II tetramers and dextramers ^11, 15^. However, appearance of CTLs was expected because of their disease-protective roles in virus infections, and indeed, CD8-deficient animals were previously shown to be highly susceptible to CVB3 infection ^80^. Nonetheless, detection of Treg cells was not expected, and their infiltration may be necessary to achieve immune homeostasis by suppressing the ongoing inflammation. Furthermore, DGE analysis revealed identical transcriptomes between CD4 and CD8 T cells, with *Ccl5, Nkg7*, and *S100a4* being prominent (**Fig 3E**), the latter two of which have been implicated in cytotoxic and fibrotic functions ^23, 35^. While expression of *Nkg7* in CD8 T cells can be related to cytotoxic functionality, its expression in CD4 T cells may suggest the possibility that CD4 T cells may function as cytotoxic CD4 T cells. We noted the upregulation of *Nkg7* in Th17 cells, as well, and this was a new finding in myocarditis (**Fig 3F**). Recent reports have shown an upregulated *Nkg7* expression in CD4^+^ T cells in patients treated for visceral leishmaniasis, and *Nkg7*^−/−^ mice infected with *Leishmania donovani* and *Plasmodium berghei* had reduced inflammation ^35^. These observations suggest that *Nkg7* expression may promote pro-inflammatory functions of Th17 cells. Consistent with this notion, transcriptome analysis in the Treg cells revealed the expression of various genes (*Nfg7, Prfn1*) that are implicated in cytotoxicity (**Fig 3F**). This observation, however, may raise the question whether Treg cells can contribute to the persistence of virus infection by killing virus-reactive CTLs or suppress autoreactive T cell responses via cytotoxicity. The former possibility is unlikely because adoptive transfer of Treg cells can suppress the development of CVB myocarditis ^81^. Nonetheless, expression of *Ly6a*, the marker of double negative Tregs, was not expected in CD8 T cells (**Fig 3F**), suggesting whether a proportion of CD8^+^ *Ly6a*^+^ T cells can function as Treg cells in viral myocarditis.

Although fibroblasts form a major part of the cardiac cellulome and are implicated in the acute phase of CVB infection (**Fig 1F, G, Table S2**), their role in post-infectious myocarditis phase has not yet been studied in detail ^82, 83^. Our scRNAseq analysis revealed 12 distinct clusters that express overlapping genes (**Fig 4A, B, C**). Several of these (*Wif1a, Npy*)^47^ have been previously shown to be involved in wound healing, ventricular/cardiac remodeling, and myocardial fibrosis in DCM pathogenesis (*Ltbp2*, *Thbs4* and *Tgfbi*) ^24, 42, 43^. However, a subset of genes (*Mt1, Mt2, Cxcl1*)^25, 30^ that have immune modulatory roles and promote angiogenesis (*Vegfd*) were also detected in various clusters. By evaluating the top eight genes in all clusters, we were able to identify two categories of subclusters with one gene in each that mediated cardiac remodeling events and anti-viral and immune activation functions (**Fig 4C**). These functionalities of fibroblasts could be further supported by evaluating the DGE of various transcripts which had roles in wound healing, regulation of inflammation, and immune responses (**Fig 4D**). Our data also revealed detection of two transcripts (*Cyb5a* and *Fxpyd6*) (**Fig 4E**) that were not previously associated with the known cardiac injury marker *Wif1a*, suggesting the possibility of their use as novel cardiac injury markers. Overall, although the composition of fibroblasts was complex, their transcriptome profiles revealed a diverse role in fibrosis, immune activation, and inflammatory functions.

Among other cells, NK cells and ILCs were low in number (**Fig 1G, Table S2**), but several genes with known functions were noted in NK cells, including *Tmem176b*, which has a role in fibrosis. Likewise, DGE analysis in the neutrophils revealed transcripts that have varied functions, such as negative regulation of inflammation, antiviral response, antigen-presentation functions, and wound healing (**Fig 6D**). Notably, *Ly6g*^+^ cluster 2 had increased expression of *Il1b* indicating that neutrophils might release *Il1b* to promote pathogenic fibrosis and cardiac remodeling, eventually leading to DCM, thus providing more evidence to previous hypotheses ^64, 84^. However, the IL-17 signaling pathway appeared to be dominantly influenced by genes expressed in neutrophils, raising a question as to their reactivity to Th17 cytokines (**Fig 6E, Fig S6B**). On one hand, Th17 cells facilitate neutrophil chemotaxis, but on the other, neutrophils do not respond to IL-17, since they lack expression of *IL-17Rc* ^85^. It may be that the genes expressed in neutrophils may modulate the IL-17 signaling pathway in non-neutrophils. Unexpectedly, B cells formed a minor fraction in the heart infiltrates, and their number was significantly low in myocarditic mice (**Fig 1G, Table S2**). Analysis of their transcriptomes also did not reveal significant differences between myocarditic and healthy mice (**Fig S7C**), suggesting that B cells appear not to have a major role in chronic myocarditis.

Constructing the intercellular communication networks within the cardiac cellulome using ligand-receptor interactions in myocarditic mice indicated strong associations between CD4 and CD8 T cells, CD8 T cells and neutrophils, myeloid cells and NK cells, and fibroblasts and SMCs and myeloid cells, among others (**Fig 7A, B**). Likewise, the number of receptor-ligand pairs and interaction strengths were relatively high between fibroblasts and CD4 and CD8 T cells, and between myeloid cells and CD4 T cells and SMCs (**Fig 7C, Fig S8B**). In a virus-free setting of autoimmune myocarditis, it has been suggested that the Th17 cells-fibroblasts-monocytes/macrophage axis may be critical for development of inflammatory cardiomyopathy ^86^, which may not be relevant to viral myocarditis. Based on our data, we noted a dependency between CD4 and CD8 T cells in their functionalities, and CD8 T cells may potentially influence the effects of other innate cells, including fibroblasts. Such a possibility can be expected with virus infections, since CD8 T cells form an important component of anti-viral responses. Furthermore, by analyzing the signaling molecules in relation to receptor-ligand interactions, we noted distinct pathways to be important in reparative processes (**Fig 7F**). Increased EGF, LCK, ncWNT, and NOTCH signaling (**Fig 7F, Fig S10**), which have been reported to be necessary for CVB3 replication and cardiac fibrosis, could indicate that these pathways could be promoting the anti-inflammatory reparative process in the myocardium, as well as cardiac remodeling in the post-infectious myocarditis phase ^66–68, 70^. OSM’s role in cardiomyocyte de-differentiation has been found to be increased in mouse models of myocardial infarction and in end-stage heart failure patients ^69^. Even though OSM plays a protective role during the acute phase of myocardial damage, its prolonged expression, along with the increased infiltration of macrophages during the chronic phase, may promote functional deterioration and loss of cardiac contractility, leading to DCM and heart failure. PDGF, which has been known to be involved in fibrosis and upregulated in CVB3 infection, showed increased signaling in the post-infectious myocarditis phase. Profibrotic *Tgfbi* was upregulated in monocyte cluster 0 and *Mfap4*^+^ in fibroblast cluster 6, and is known to promote PDGF signaling, thus suggesting a role for it in the cardiac fibrosis and remodeling phase of post-infectious myocarditis. Overall, our data indicate that T cells, myeloid cells, and fibroblasts may mainly contribute to the progression of viral myocarditis in chronically infected mice, and that the upregulated signaling pathways could be targeted for therapeutic purposes in DCM.

Finally, in our efforts to identify the potential TFs that might regulate various cellular functions, we identified *Ets1* in T cells and *Irf5,* and *Mafb* in myeloid cells which have varied roles (inflammation, differentiation of monocytes/macrophages and cardiac morphogenesis) as potential target candidates TFs (**Fig 8A**). In an *AngII* mouse model of cardiac fibrosis, a recent report showed that deletion of *Ets1* from ECs reduced cardiac fibrosis and hypertrophy^87^. Upregulated expression of *Ets1* in our data, which was regulating expression of target genes implicated in cardiac fibrosis and heart failure may mean that targeting this TF could be beneficial in attenuating the transition to DCM. Although the roles for *Irf5* and *Mafb* have not been defined clearly in myocarditis, our data indicates their important role in controlling expression of M2-specific genes and transcripts involved in cardiac fibrosis/remodeling/heart failure during the post-infectious phase of myocarditis. Since, these TFs can act as activators or repressors of various genes, determination of the myocarditis phenotype in mice deficient for TFs may permit us to evaluate their roles in viral myocarditis and develop strategies for therapeutic interventions.

In summary, we have described the cellular compositions and their transcriptome profiles in heart infiltrates using the mouse model of viral myocarditis. While T cells, myeloid cells, and fibroblasts formed a major component, the proportion of B cells was low. Although CVB3 infection has been extensively used to understand the pathogenesis of myocarditis, a long-standing question remains as to the underlying mechanisms of chronic myocarditis, with autoimmune theory as one possibility. Reports indicate detection of autoantibodies in CVB3 myocarditis, but their pathogenic role remains inclusive. We have been investigating the role of autoreactive T cells and, using MHC class II dextramers, have reported the appearance of pathogenic autoreactive T cells with specificities for multiple antigens ^11, 15^. Our scRNAseq data also point to a role for T cells, but in-depth analysis of their role can be investigated in antigen-specific (dextramer^+^) T cells by scRNAseq analysis. Likewise, detection of M2 cells was not surprising because of their role in reparative functions, and detection of fibroblasts was also expected due to their role in the formation of fibrosis. In adjuvant-induced myocarditis, studies have recently reported identification of neutrophils, macrophages, γδ T cells, Th17 cells, and Tregs with different transcriptome signatures as compared to our study with viral myocarditis ^88^. These variations are expected because of fundamental differences between the two (virus-free, adjuvant vs virus). Of note, we could not study the compositions of non-immune cells of importance in the myocardium, which include cardiomyocytes, SMCs, pericytes, and ECs. scRNAseq analysis of their purified populations using single-nuclei RNA seq may yield new insights into viral pathogenesis, and we will investigate this in the future. For example, although low in number, we investigated the transcriptome profile of ECs, leading us to note upregulation of TFs, which have a role in apoptosis of ECs and inflammation (*Atf3*, *Irf1*, and *Stat6*), with a corresponding downregulation of those involved in EC survival and angiogenesis (*Myc* and *Nfe212*) in myocarditic animals as compared to controls (data not shown). Overall, our scRNAseq analysis offers a new dimension to understanding the post-infectious phase of viral myocarditis and associated pathogenic cardiac remodeling. Detailed investigation of these novel genes, markers, TFs, and signaling pathways may offer new therapeutic targets of clinical relevance for DCM and heart failure.

## Materials and Methods

### Mice

Six-to-eight-week-old, and nine-11-week-old male A/J mice (H-2^a^) were procured from Jackson Laboratory (Bar Harbor, ME) and maintained according to the Institutional Animal Care and Use Committee’s guidelines of the University of Nebraska-Lincoln (protocol #: 1904), Lincoln, NE. Infection studies were performed based on biosafety level 2 guidelines. When animals were found to have persistent clinical signs, such as failure to move when physically touched or prodded, or failure to eat or drink, they were euthanized using a carbon dioxide chamber as recommended by the Panel on Euthanasia of the American Veterinary Medical Association.

### Virus propagation and infection

The Nancy strain of CVB3 was procured from the American Type Culture Collection (ATCC, Manassas, VA, USA), and the virus was titrated in Vero cells (ATCC). The adherent Vero cells were grown to 80 to 90% confluence in 75cm^2^ flasks in EMEM/10% fetal bovine serum (FBS) and were later infected with CVB3 with multiplicity of infection 1 in EMEM containing no FBS. After incubation at 37⁰ C for 1 hour with gentle intermittent rocking, maintenance medium (EMEM/2% FBS) was added. Based on the cytopathic effect of virus during the next 1 to 2 days, supernatants containing virus were harvested. After determining 50% tissue culture infective dose (TCID_50_) values based on the Reed-Muench method, the virus stocks were aliquoted and preserved at -80⁰ C. To infect mice, virus stock diluted in 1x PBS to contain 10,000 TCID_50_ in 200 µl was administered intraperitoneally (i.p.). Animals were monitored closely, cages were changed once in 2 days, and body weights were taken daily until termination. In addition, an alternative food and fluid source, trans gel diet (ClearH2O, Portland, ME, USA), was placed on the cage floor as needed.

### Heart single-cell preparation

Single-cell suspensions from mouse hearts were prepared as previously described (Pinto 2013, 2016). Briefly, male CVB3-infected mice and their age-matched healthy control mice were euthanized on day 21 post-infection using 2% CO_2_ in an asphyxiation chamber as per the IACUC guidelines. The hearts were perfused using perfusion buffer (1 × DPBS with 0.8 mM CaCl2, 5 ml/min for 5 minutes) until the liver was completely blanched and appeared pale yellow/brown in color. Next, hearts were isolated, their atria and valves were removed, and the whole heart was minced to ∼0.5-1 mm cubes using surgical scissors. Minced heart tissue was digested in 3 mL of digestion buffer (2 mg/mL Collagenase IV [Worthington Biochemical], 1.2 units/mL Dispase II [Sigma-Aldrich], in perfusion buffer) for ∼45 minutes at 37° C using a rotating holder, with tissue suspension triturated once in every 15 minutes with 1000 μl wide-bore micropipette tips. The cell suspension was filtered through a 70 μm nylon filter mesh to remove any residual undigested tissue pieces. The filtrate was then diluted in ∼15 mL perfusion buffer, and the cells were pelleted at ∼200 × G for 20 minutes with no centrifuge brakes engaged. Cell supernatant was then aspirated, and the pellet was re-suspended in ∼15 mL of 1 × HBSS (Sigma-Aldrich) + 0.8 mM CaCl2. The cells were pelleted again as described above. In order to remove unwanted debris, a debris removal kit was used as per the manufacturer’s guidelines (Miltenyi Biotec, San Diego, CA). The final debris-free cell pellet was resuspended in 1000 μL of 2% FBS in RPMI medium for downstream cell sorting using flow cytometry.

### Flow cytometry and sorting

Freshly prepared cardiac single-cell suspensions from healthy and myocarditic hearts were subjected to surface staining with Annexin V (Biolegend, San Diego, CA) and propidium iodide (PI; Biolegend). In brief, cells were first washed with Annexin V binding buffer, followed by surface staining with Annexin V (1:200 vol/vol) and PI (1:100, vol/vol) at room temperature (25°C) for 15 minutes in the dark. Cells were then resuspended and sorted by flow cytometry (FACSAria II, BD Biosciences, San Jose, CA). Only singlets that were viable and non-apoptotic (Annexin V^-^ PI^-^) were sorted and collected in tubes containing RPMI with 2% FBS.

### Sample processing and sequencing

Two replicates, with n=7 mice per treatment group, were used for heart sample processing. Approximately 16,000 cells were loaded onto a single channel of the 10X Genomics chromium controller (10X Genomics, Pleasanton, CA), with a target recovery of ∼10,000 cells using the chromium v2 and v3 single-cell reagent kit. After the generation of single-cell gel bead-in-emulsions, cDNA was synthesized using a C1000 Touch Thermal Cycler (Bio-Rad Laboratories, Hercules, CA) and amplified for 11 cycles as per the manufacturer’s protocol. Quality control (QC) and quantification were performed using the Agilent 2100 bioanalyzer (Agilent Technologies, Santa Clara, CA) as per the manufacturer’s guidelines. Amplified cDNA (50 ng) was used to construct 3’ expression libraries, and the libraries were pooled and run on an Illumina HiSeq 4000. Each lane consisted of 150 base-pair, paired-end reads. The Illumina basecall files were converted to FASTQ format, and these files were aligned to the murine genome (mm10) using the CellRanger v3.0.2 pipeline as described by the manufacturer. Across aligned cells, the mean number of reads per cell was 39,923, with an average of 95.3% of reads mapped to the mm10 genome.

### Single-cell data processing and analysis

Initial processing of cells isolated from the heart in myocarditis run 1 (n=2,617), myocarditis run 2 (n=10,618), control run1 (n=1,528), and control run 2 (n=8,201) was performed using the Seurat R package (v3.0.2) ^16, 89^. Samples were normalized using the sctransform approach ^90^ with default settings. The transformed data was then formed into a single data set using canonical correlational analysis and mutual nearest neighbors (MNN) as described by Stuart et al ^89^. Dimensional reduction to form the uniform manifold approximation and project (UMAP) utilized the top 30 calculated dimensions and a resolution of 0.6. The schex R package (v1.1.5) was used to visualize mRNA expression of lineage-specific or highly differential markers by converting the UMAP manifold into hexbin quantifications of the proportion of single cells with the indicated gene expressed. Default binning was set at 80, unless otherwise indicated in the figure legend.

Cell type identification utilized the SingleR (v1.0.1) R package ^18^ with correlations of the single-cell expression values with transcriptional profiles from pure cell populations in the Immgen ^17^. In addition to correlations, canonical markers for cell lineages were utilized and are available in Table S1. Differential gene expression utilized the Wilcoxon rank sum test on count-level mRNA data. For differential gene expression across clusters or subclusters, the *FindAllMarkers* function in the Seurat package was used, employing the log-fold change threshold > 0.25, minimum group percentage = 10%, and pseudocount = 0.1. Differential comparisons between conditions utilized the *FindMarkers* function in Seurat without filtering and a pseudocount = 0.1. Multiple hypothesis correction was reported using the Bonferroni method. Cell cycle regression was performed in Seurat using the *CellCycleScoring* function and genes derived from Nestorowa et al ^91^. Genes were isolated by calling *cc.genes.updated.2019* in R and then converting into murine nomenclature. Gene set enrichment analysis was performed using the escape R package (v0.99.0). Differential enrichment analysis was performed using the *getSignificance* function in escape that is based on the limma R package linear fit model.

### Cell trajectory analysis

Cell trajectory analysis used the Slingshot (v1.6.0) R package with default settings for the slingshot function and using the UMAP embeddings from the subclustering for each cell type. Ranked importance of genes was calculated using the top 300 variable genes, and rsample (v0.0.9) and tidymodels (v0.1.0) R packages were used to generate random forest models based on a training data set of 75% of the cells. The *rand_forest* function in the parsnip (v0.1.1) R package was used, with mtry set to 200, trees to 1400, and minimum number of data points in a node equal to 15 across all cell types.

### GO and pathway enrichment analysis of DEGs

GO pathway enrichment analysis of myocarditis-related DEGs was performed by Metascape (http://metascape.org/gp/index.html) (version 3.5). Results were visualized using the ggplot2 R package (version 3.2.1). Single-cell normalized enrichment scores were calculated using the escape (v1.0.1) R package ^92^. From this analysis, differentially expressed ligand and receptor between myocarditic and healthy controls for indicated cell types were extracted to use for the size/count of the dot plot. Differential gene set enrichment utilized the Welch’s T test with the Bonferroni adjustment for multiple hypothesis correction comparing individual cells in myocarditis versus healthy controls.

### Intercellular communication analysis

Cell-cell interactions based on the expression of known ligand-receptor pairs in different cell types were inferred using CellChat (version 1.0.0) R package^65^. We used the default settings to predict major signaling interactions of cells and how these cells and signals coordinate various functions. In brief, we followed the workflow recommended in CellChat and loaded the normalized counts into CellChat and applied the preprocessing functions *identifyOverExpressedGenes*, *identifyOverExpressedInteractions*, and *projectData* with default parameters set. For the analysis of ligand-receptor interactions, the functions *computeCommunProb*, *computeCommunProbPathway*, and *aggregateNet* were applied using default parameters. Finally, we classified signaling pathways and depicted conserved and context-specific pathways between myocarditis and healthy hearts.

### Analysis of TF regulatory network

TF regulatory network analysis was performed using SCENIC ^71^ (version 1.1.2.2) with default parameters and a co-expression method set to top 50 results per target. Murine mm9 TFs were downloaded using RcisTarget (version 1.6.0) as a reference. Enriched TF-binding motifs predicted candidate target genes (regulons), and regulon activity was inferred by RcisTarget. Resulting AUC enrichments for individual cells were attached to the Seurat object, and the median by cluster and condition were visualized using the pheatmap R package (v1.0.12). Transcription factor regulons were concatenated across the results from the motif enrichment step and graphed using the igraph (v1.2.6) R package.

### Statistical analyses

Statistical Analyses were performed in R (v3.6.3). Two-sample significance testing utilized Welch’s T test, with significance testing for more than three samples utilizing one-way analysis of variance (ANOVA) with Tukey honest significance determination for correcting multiple comparisons. Two-proportion Z-tests were performed using the total number of cells in each condition as the number of trials and without a prior for proportion.

## Author contributions

N.L. and J.R. conceptualized the study; N.L. and R.A. performed the experiments; N.L., N.B., and J.R. processed, analyzed, and interpreted the data; N.L. and J.R. drafted the manuscript; N.L., N.B., T.K.S., and J.R. edited the manuscript.

## Acknowledgments

This work was supported by the Transformational grant from the American Heart Association and the institutional funds from the University of Nebraska-Lincoln to J.R.

## Competing interests

The authors declare no competing interests.

## Data availability

Raw sequencing data and quantified gene expression counts for single-cell RNA sequencing are available at the Gene Expression Omnibus at GSE174458. Any other data relevant to this study are available from the authors upon reasonable request.

## Supplementary Information

Fig S1. **Schematic representation for sorting single cells from heart infiltrates.** Groups of mice were infected with or without CVB3, and after 21 days, hearts were collected at euthanasia following perfusion. Hearts were enzymatically digested to obtain single cell suspensions, and cells were then stained with annexin-V and PI for sorting by flow cytometry, where viable (annexin-V-, PI-) cells were sorted by gating the singlets.

Fig S2. **Canonical markers and GSEA in myeloid cells. A.** UMAPs showing percentage expression of indicated markers in myeloid cell clusters. **B.** Enriched GO terms with respect to pathways upregulated in myeloid cells of myocarditic mice. **C.** Enriched pathways upregulated in myeloid cell clusters 0, 1, 3 of myocarditic mice.

Fig S3. **Differentially expressed genes and their pathway analysis in T cells. A.** UMAPs showing the percentage expression of canonical T cell markers. **B.** Cell cycle phases by clusters in myocarditic mice in relation to healthy group. **C.** Top 15 genes upregulated and downregulated in myocarditis versus healthy controls by log-fold change in myocarditic mice. **D.** Pathway analysis corresponding to Th17 subset (cluster 1) and CTLs (cluster 2).

Fig S4. **Cell cycle phases and GSEA of fibroblast cell clusters in heart infiltrates. A.** Cell cycle phases by clusters between groups. **B**. Pathway analysis is shown for clusters 1, 3, 5, and 6.

Fig S5. **Differentially expressed genes and their pathway analysis in NK cells. A.** Top 15 upregulated and downregulated genes in myocarditic mice versus healthy controls by log-fold change in NK cells. **B**. Differential expression of the top 10 genes in *Gzma*^+^ cluster 0, *Tcf7*^+^ cluster 1 of myocarditic mice relative to healthy group. **C.** Pathway analysis for *Gzma*^+^ cluster 0.

Fig S6. **Differentially expressed genes and their pathway analysis in neutrophils. A.** Overall differential expression of genes in neutrophils is shown in the volcano plot (left), including the top 10 genes in the bar plot (right panel). **B.** Pathway analysis for clusters 0, 1, and 2.

Fig S7. **Analysis of B cell clusters in myocarditic hearts A.** UMAP showing different clusters of B cells in control and myocarditic mice. **B.** Distribution of B cells and their relative proportions in control and myocarditic mice. **C.** Volcano plot showing the differentially expressed genes.

Fig S8. **Intercellular communication in the healthy and myocarditic cardiac cellulome. A.** Circle plot showing intercellular communication between major cardiac cell types for control hearts using CellChat R workflow. The lines originating from a cell type indicate ligands being broadcast, with these lines connecting to the cell type where the receptors are expressed. Thickness of the line is proportional to the number of unique ligand-receptor interactions, with loops representing autocrine circuits. **B**. Heatmap of differential interaction strength between myocarditic and healthy mice in the cell-cell communication network. The top-colored bar plot indicates the sum of column values (incoming signaling), and the right bar plot indicates the sum of row values (outgoing signaling). Red indicates increased signaling in myocarditis, and blue indicates decreased signaling.

Fig S9. **Intercellular interactions between cell types in control and myocarditic hearts.** A detailed view of ligand and cognate receptor interaction for each cell type in myocarditic (**A**), and control (**B**) hearts.

Fig S10. **Major signaling pathways inferred from intercellular communication in the myocarditic hearts.** Signaling interactions for specific pathways between cells in the cardiac cellulome in myocarditic mice. Signals are being sent from the source (senders) along the *y* axis to the targets (receivers) along the *x* axis.

Fig S11. **GO functions of TF-regulated target genes.** The upregulated GO functions of various TF-regulated target genes found enriched in **(A)** T cells, (**B**) myeloid cells, (**C**) neutrophils, and (**D**) NK cells.

Fig S12. **Analysis of transcription factors in neutrophils, NK cells and fibroblasts.** Enrichment of TFs in neutrophils, NK cells and fibroblasts from control and myocarditic mice as analyzed by SCENIC.

## Notes

### Competing Interest Statement

The authors have declared no competing interest.

