## Supplementary material for "Dissecting the Cellular Landscape and Transcriptome Network in Viral Myocarditis by Single-Cell RNA Sequencing": Final Supplementary figures.pdf

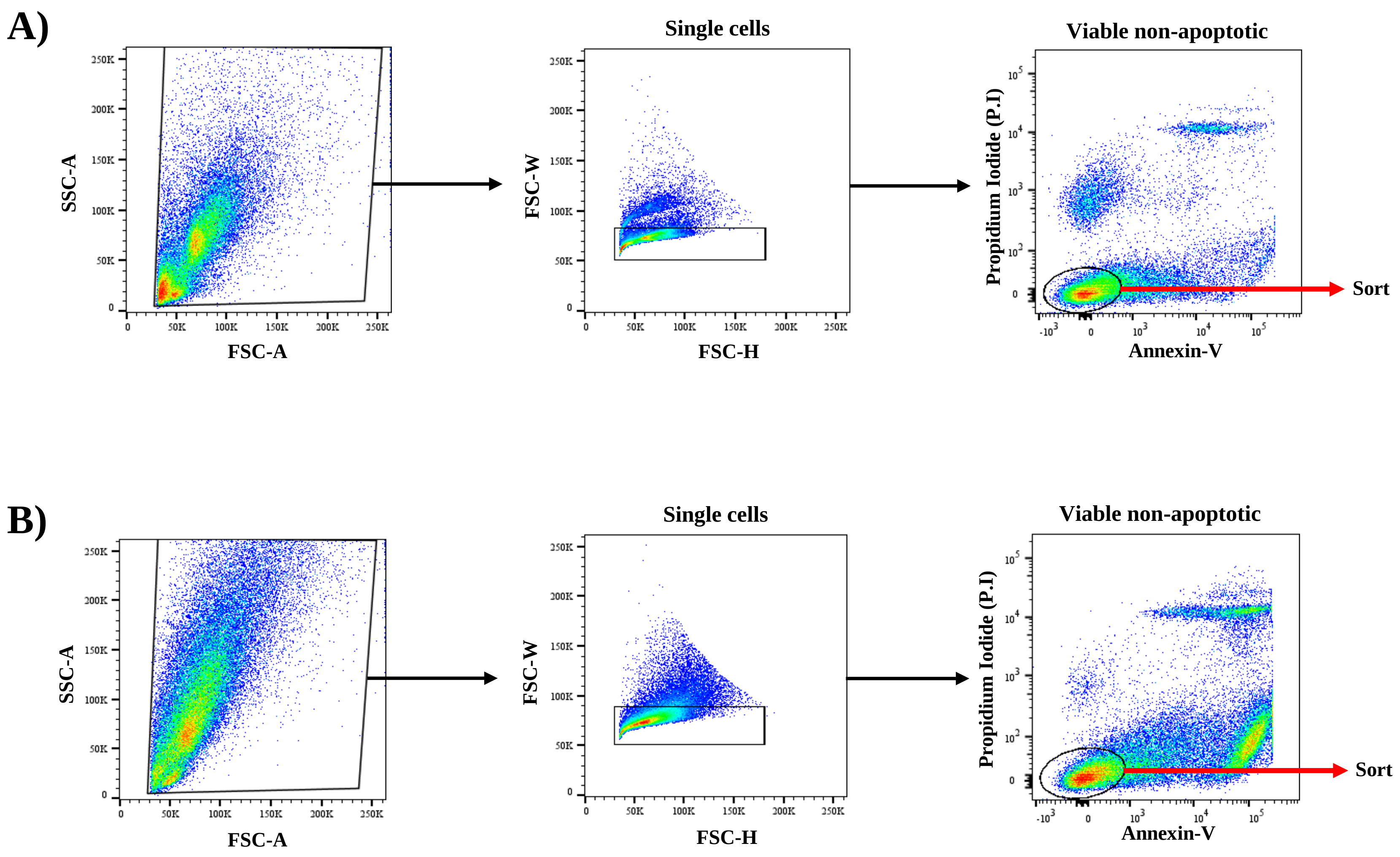


**Fig S1: Schematic represntation for sorting single cells from heart infiltrates.**

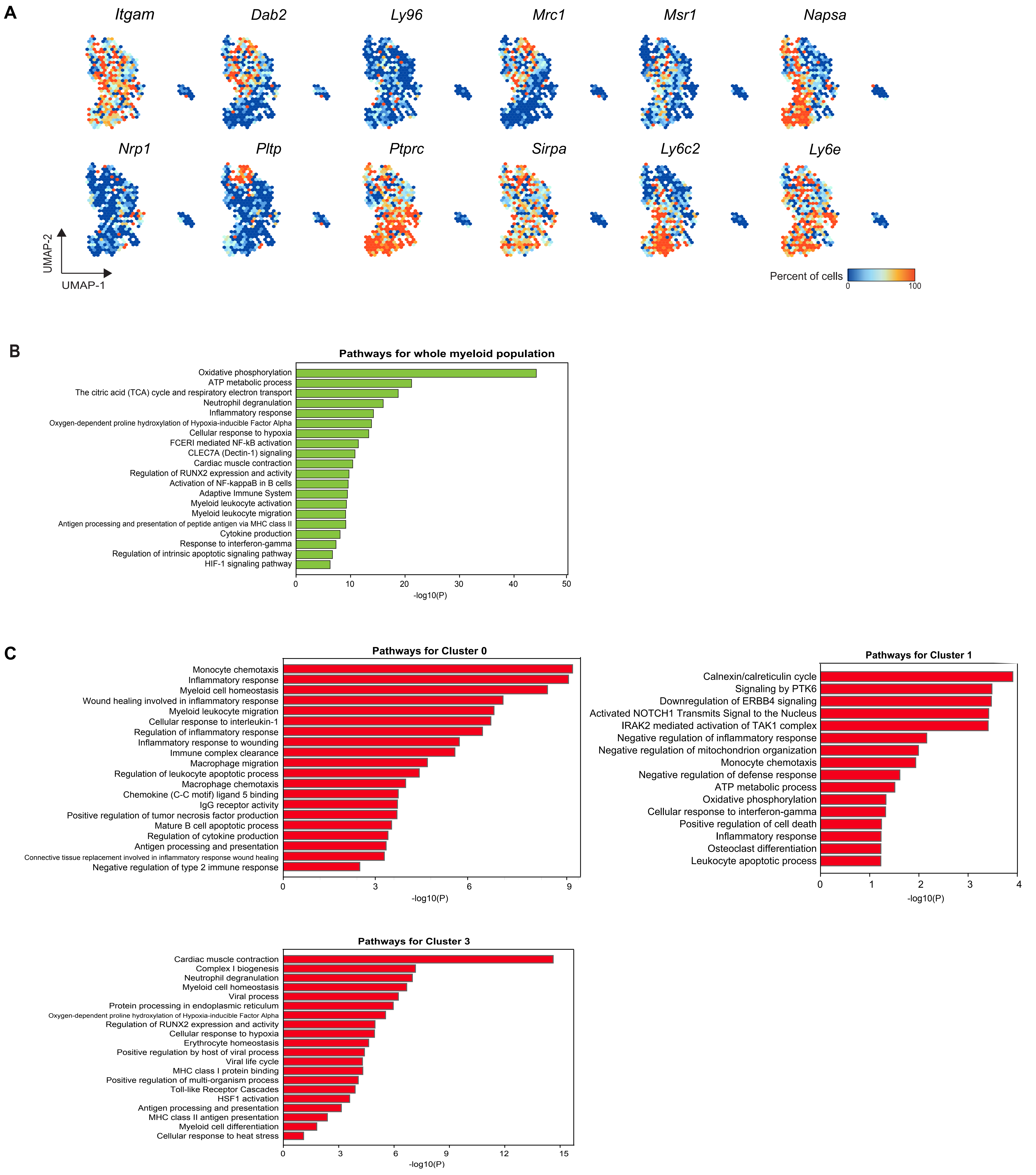


**Fig S2: Canonical markers and GSEA in myeloid cells**

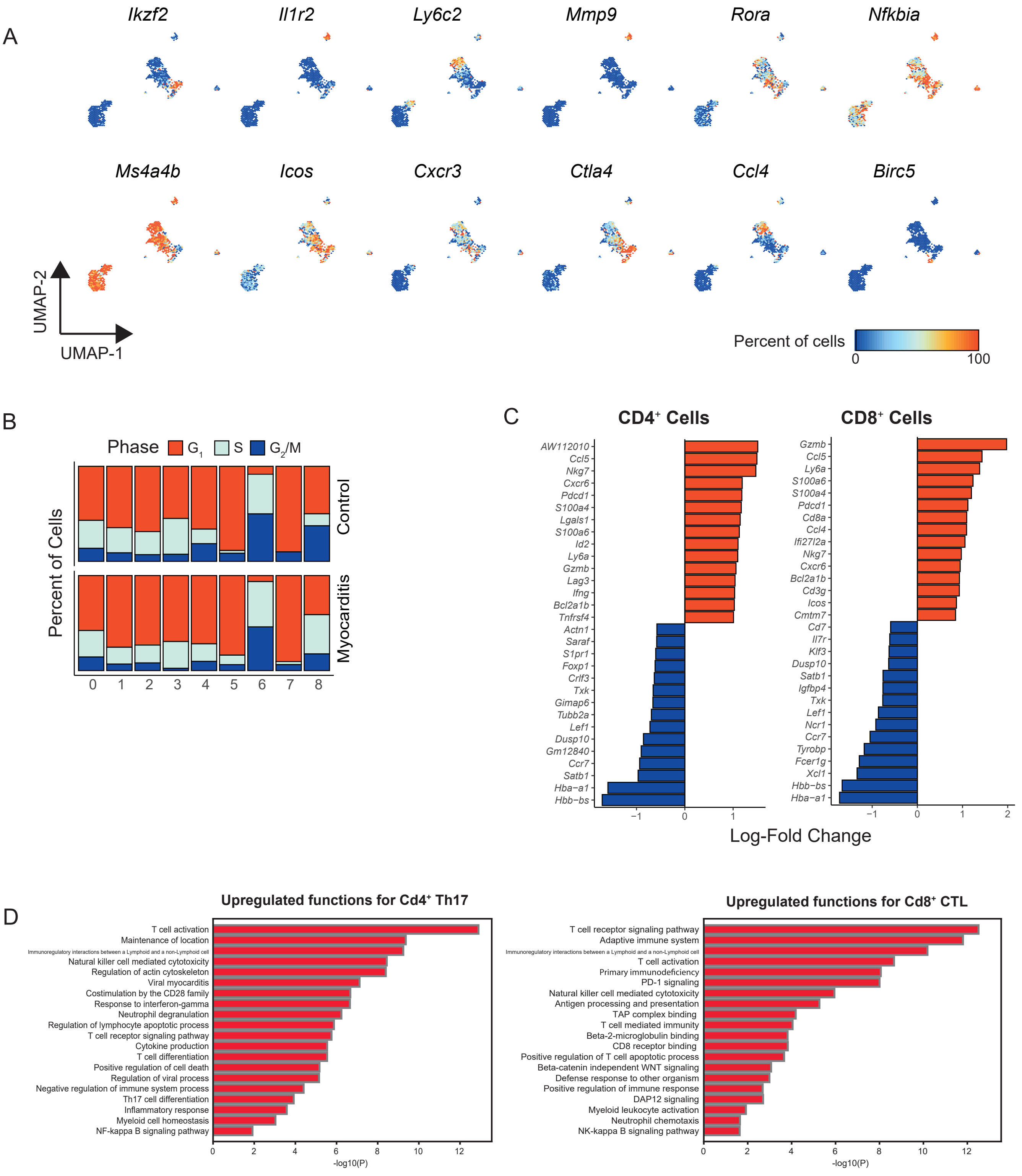


**Fig S3: Differentially expressed genes and their pathway analysis in T cells.**

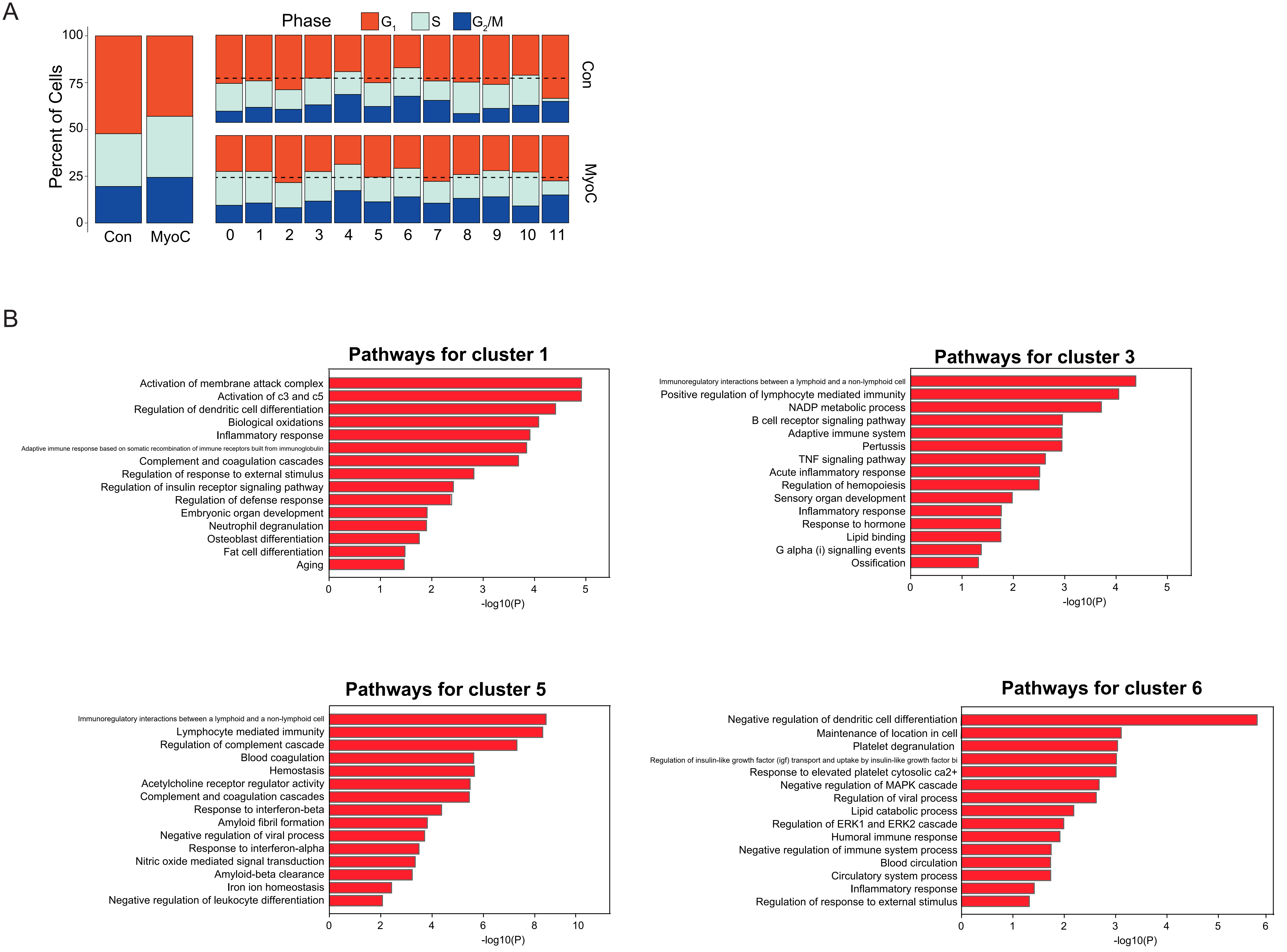


**Fig S4: Cell cycle phases and GSEA of fibroblast cell clusters in heart infiltrates.**

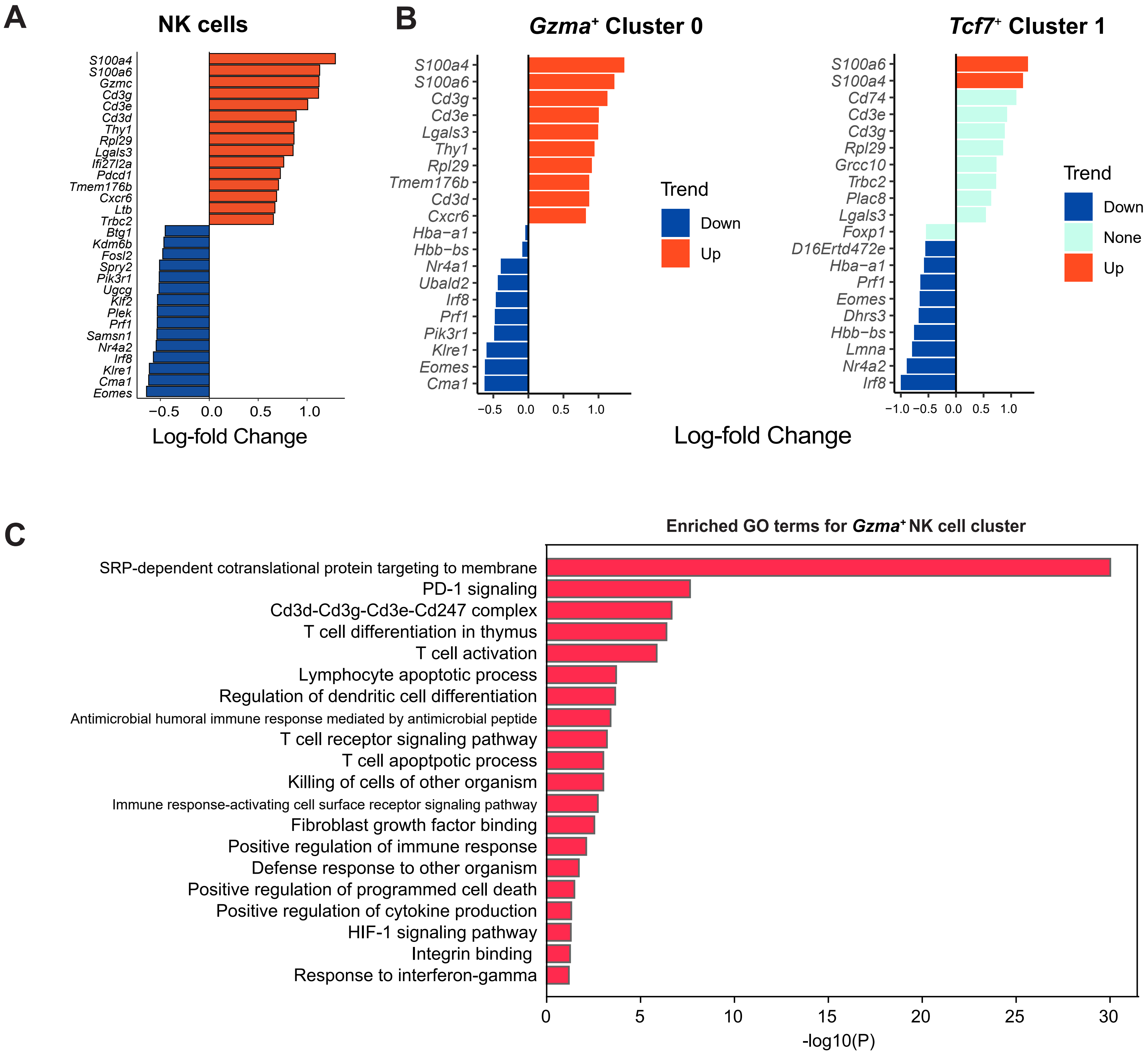


**Fig S5: Differentially expressed genes and their pathway analysis in NK cells**

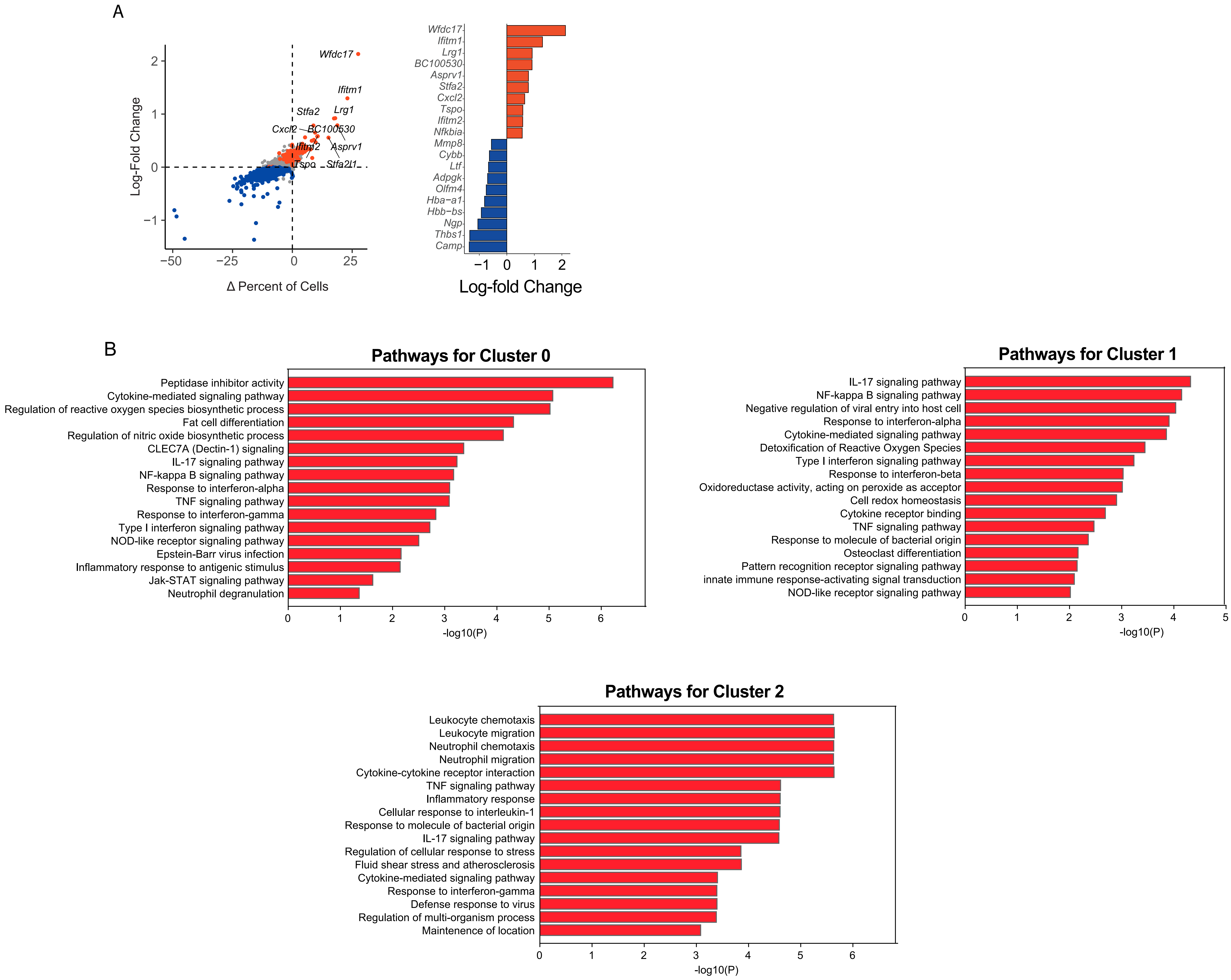


**Fig S6: Differentially expressed genes and their pathway analysis in neutrophils.**

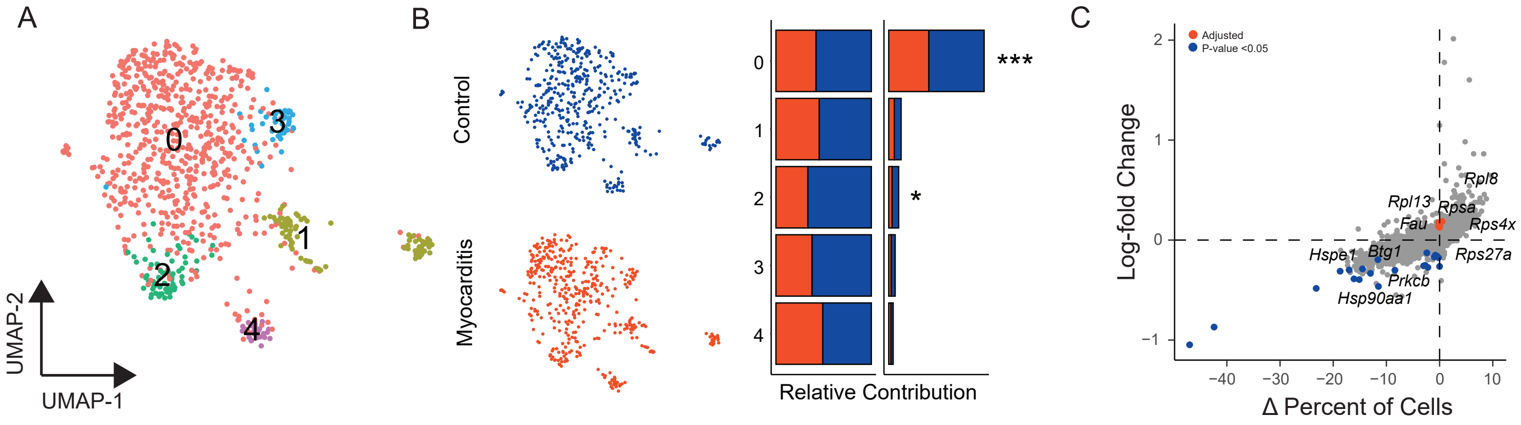


**Fig S7: Analysis of B cell clusters in heart infiltrates.**

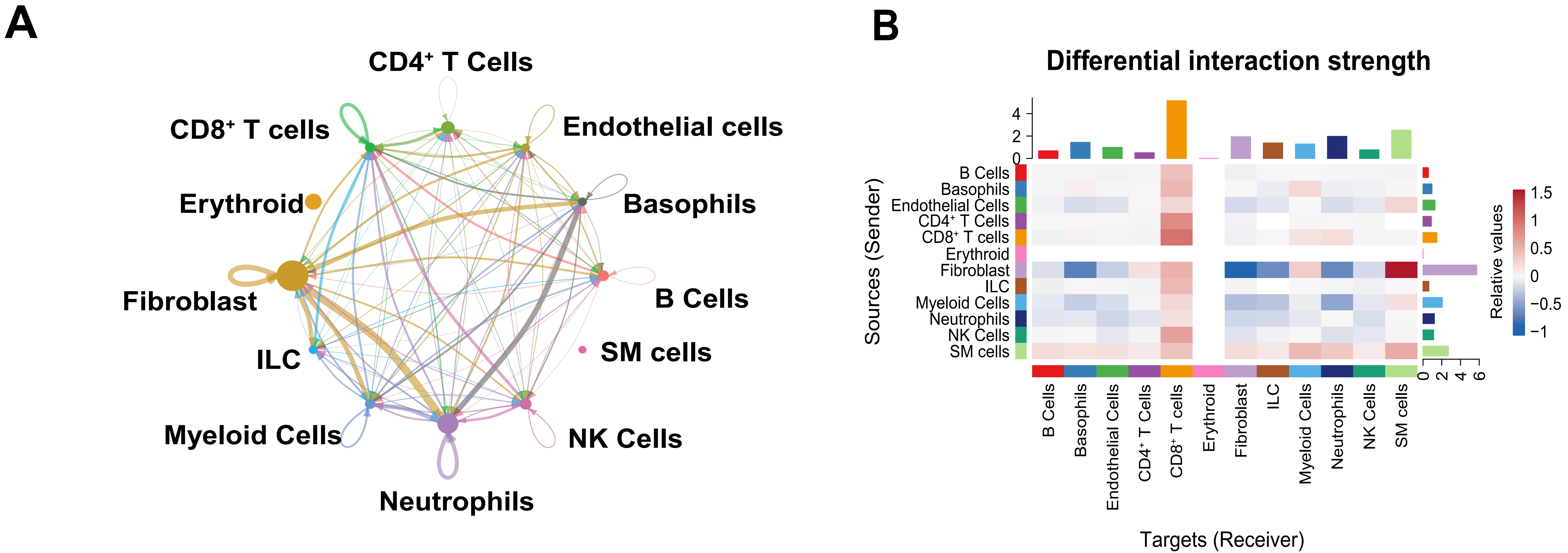


**Fig S8: Intercellular communication in the healthy and myocarditic cardiac cellulome**

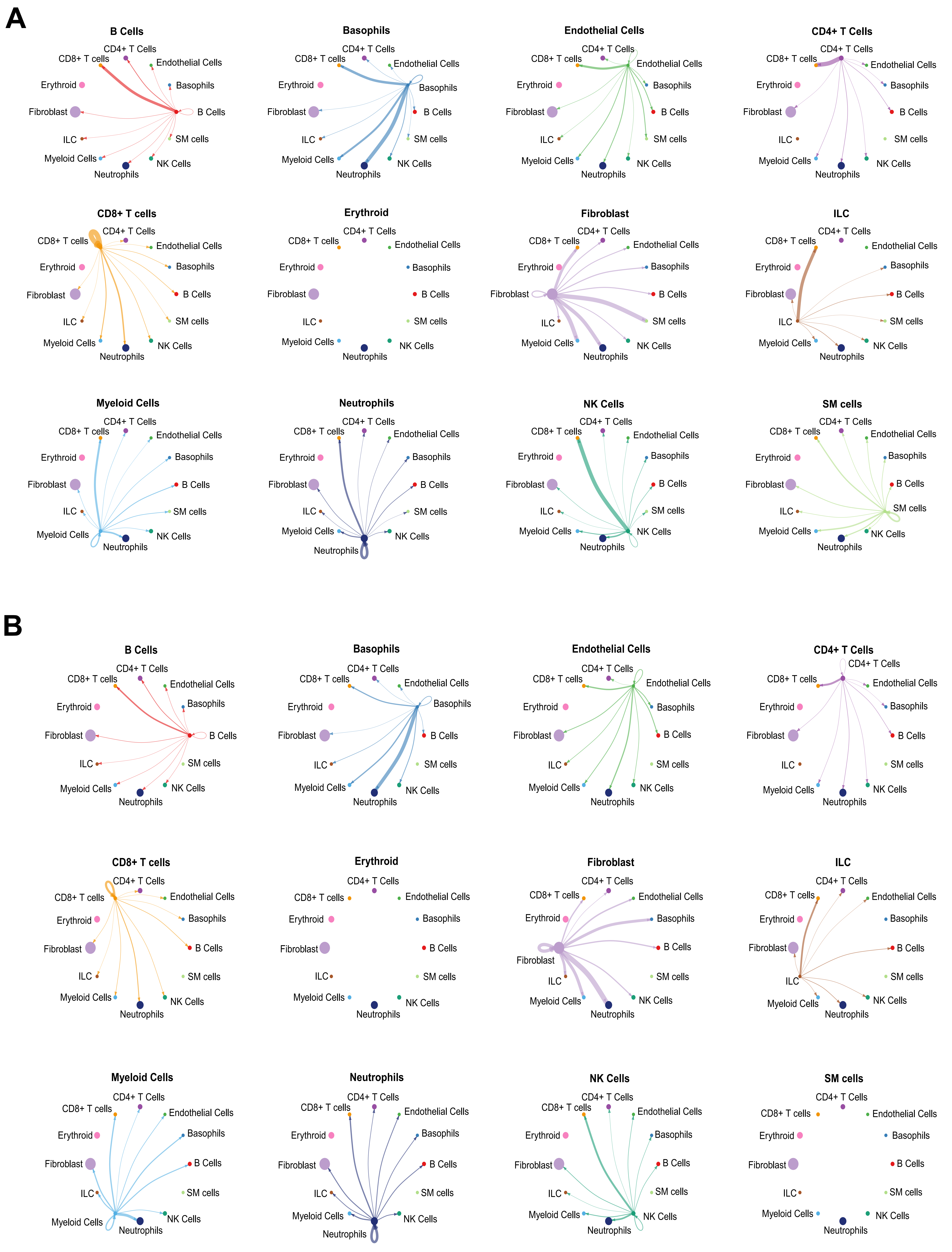


**Fig S9: Intercellular interactions between cell types in control and myocarditic hearts**

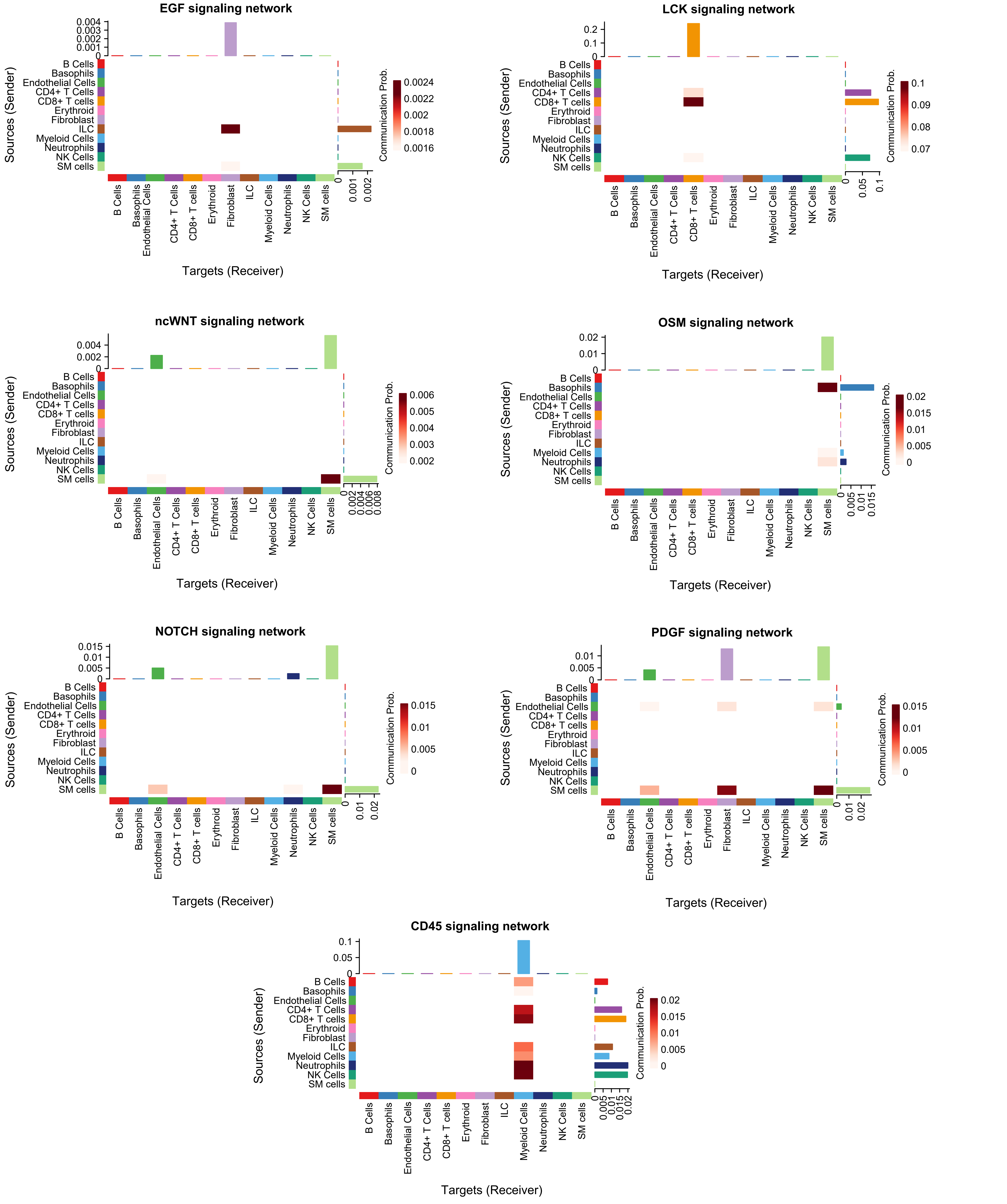


**Fig S10: Major signaling pathways inferred from intercellular communication in the myocarditic hearts**

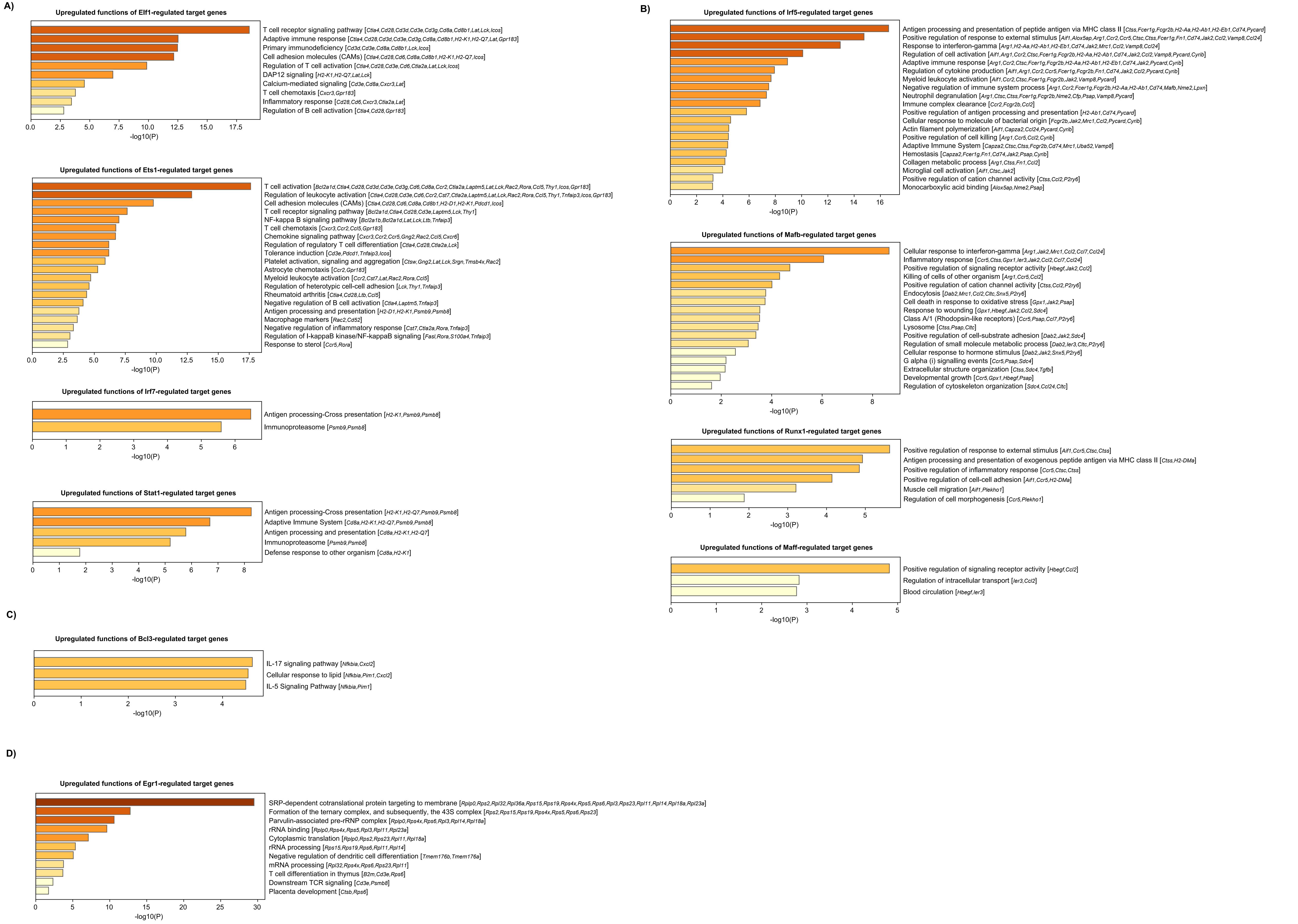


**Fig S11: GO functions of TF-regulated target genes**

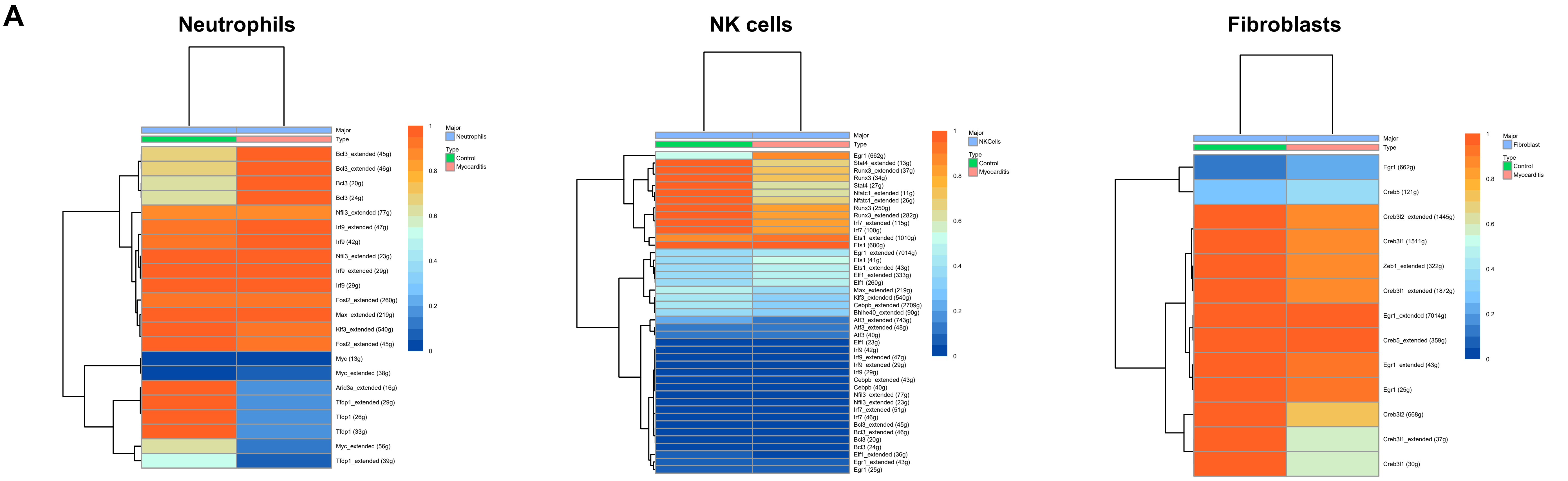


**Fig S12: Analysis of transcription factors in neutrophils, NK cells and fibroblasts**
