## Supplementary material for "Dissecting the Cellular Landscape and Transcriptome Network in Viral Myocarditis by Single-Cell RNA Sequencing": Table S1.docx

| **Cell Types** | **Marker genes** |
| --- | --- |
| B cells | *Cd19, Ighm, Bank1, Cd79b, Fcmr, H2-Ob, Ms4a1, Tnfrsf13c* |
| Basophils | *Mcpt4, Mcpt8, Ifitm1, Cyp11a1* |
| Endothelial Cells | *Cdh5, Egfl7, Apold1, Arhgef15, Emcn, Gpihbp1, Rasgrp3, Rbp7, Tie1* |
| T Cells | *Cd3e, Cd3g, Cd4, Cd8a, Cd8b1, Lat, Tcf7, Trbc1, Trbc2, Ctla4, Tnfrsf4* |
| Erythroid cells | *Hba-a1, Hba-a2, Hbb-bt, Alas2, Bpgm, Snca* |
| Fibroblast | *Serpinh1, Pcsk6, Col1a2, Col5a1, Lamb1, Tcf21, Lpl* |
| Innate lymphoid cells | *Gata3, Rorc, Tbx21* |
| Myeloid Cells | *Ccr2, Cd209a, Adreg1, Msr1, Chil3, Arg1, Ms4a6d, Ms4a6c* |
| Neutrophils | *Hp, Ccrl2, Lcn2, Pglyrp1, Cxcl2, Ngp* |
| Natural killer Cells | *Ncr1, Gzma, Gzmb, Eomes, Klre1, Klrk1, Ncr1, Prf1* |
| Smooth muscle cells | *Myh11, Mylk, Lmod1, Mustn1, Nrip2, Cnn1, Des, Grip2, Kcnmb1* |

**Table S1: Marker genes used for cell assignments**
