## Supplementary material for "Dissecting the Cellular Landscape and Transcriptome Network in Viral Myocarditis by Single-Cell RNA Sequencing": Table S2.docx

| **Cell Types** | **Control** | **Myocarditis** |
| --- | --- | --- |
| B cells | 432 | 420 |
| Basophils | 24 | 26 |
| Endothelial Cells | 13 | 47 |
| CD4^+^ T Cells | 808 | 1369 |
| CD8^+^ T cells | 266 | 586 |
| Erythroid cells | 1448 | 647 |
| Fibroblast | 3828 | 6258 |
| Innate lymphoid cells | 55 | 42 |
| Myeloid Cells | 305 | 1061 |
| Neutrophils | 2030 | 2471 |
| Natural killer Cells | 524 | 295 |
| Smooth muscle cells | 1 | 29 |

**Table S2: Total cell numbers in different cell types after QC in myocarditis and control samples**
